# Modelling the distribution of rare and data-poor diadromous fish at sea for protected area management

**DOI:** 10.1101/2022.10.24.513530

**Authors:** Sophie A. M. Elliott, Anthony Acou, Laurent Beaulaton, Jérôme Guitton, Elodie Réveillac, Etienne Rivot

## Abstract

Anthropogenic pressures have resulted in declines in diadromous fish. Many diadromous fish which were commercially important are now threatened and protected. Little is known about their marine life history phases, and no observation-based Species Distribution Model exists for this group of species at sea. Yet, fisheries dependent and independent data could provide new insights into the distribution of diadromous fish at sea.

We collated a database of 168 904 hauls from fisheries observer bycatch data and scientific fisheries surveys, from eastern Atlantic and Mediterranean waters. The distribution of eleven rare and data-poor diadromous fish (shads, lampreys, salmonids, the European eel, the thinlip mullet, smelt and the European flounder) were modelled. A Bayesian site occupancy model, that incorporates imperfect detection to account for repeat detections and non-detections, the non-random nature of fishing gear type and spatial autocorrelation was used. From the model outputs, we explored bycatch risk and the role of MPAs, required under the Marine Strategy Framework Directive and Habitat Directive and assessed.

Diadromous fish were observed within relatively shallow coastal areas. Species specific gear bycatch trends were observed. Core distribution areas corresponded to their known water basin presence, indicating connectivity with their freshwater habitats. Numerous Habitat Directive Marine Protected Areas were found to be of relevance.

Given the coastal distribution of these species, they are exposed to higher anthropogenic pressures from both terrestrial and marine environments. Risk of bycatch at sea for most species appears to be low. Nonetheless, for threatened individuals, even a small amount of bycatch may impact their populations, especially since misreporting is likely to be high. Differences in catchability between gears highlight potential benefits of limiting access of certain gears within protected areas to reduce bycatch.

## 1. Introduction

Diadromous fish undertake long distance migrations between freshwater and marine ecosystems during their varying life history stages (Lassalle et al., 2008; Legrand et al., 2020; Limburg and Waldman, 2009; McDowall, 2009). They are particularly vulnerable since they are subject to both terrestrial and freshwater pressures (e.g., terrestrial run-off, pollution, habitat destruction, barriers to migration, climate change, fishing, etc.) (Costa et al., 2021; Limburg and Waldman, 2009; Merg et al., 2020; Verhelst et al., 2021). Furthermore, unlike many marine fish that are geographically widespread, numerous anadromous fish (e.g., shad and salmonids) form river-specific populations that are more susceptible to extinction (Limburg and Waldman, 2009; McDowall, 2009). As a result, most diadromous species native to the northern Atlantic Ocean have declined in abundance by at least 90% since the end of the late 19^th^ century (Drouineau et al., 2018; Limburg and Waldman, 2009; Waldman and Quinn, 2022).

To try and halt the loss of diadromous fish populations, various International Agreements (e.g., Bern Convention, Convention of Migratory Species) and legislations (e.g., EU Habitat Directive, EU Marine Strategy Framework Directive (MSFD)) have been enacted (Table 1). Measures implemented focus principally on freshwater or estuarine habitats, such as restoration of river continuity as required by the EU Water Framework Directive (WFD, 2000/60/EC) and freshwater Spatial Areas of Conservation under the Habitat Directive (HD, 92/43/EEC). However, most diadromous species remain threatened at a national level (www.nationalredlist.org; Table 1), and the improvements have yet to be attained throughout their range (Verhelst et al., 2021; Waldman and Quinn, 2022; Wilson and Veneranta, 2019). The marine life history stages and at-sea distribution of diadromous fish, largely remain a black box (Elliott et al., 2021), causing difficulty in evaluating their conservation status required under the various legislations that protect these species (Table 1; Wilson and Veneranta, 2019). Furthermore, few Marine Protected Areas (MPAs) have been designated to protect them (e.g., https://mpa.ospar.org; https://natura2000.eea.europa.eu), and at-sea bycatch is not well understood, even if it is suspected that it could contribute to significant mortality (Kappel, 2005; Stratoudakis et al., 2016; Verhelst et al., 2021; Wilson and Veneranta, 2019).

**Table 1.**
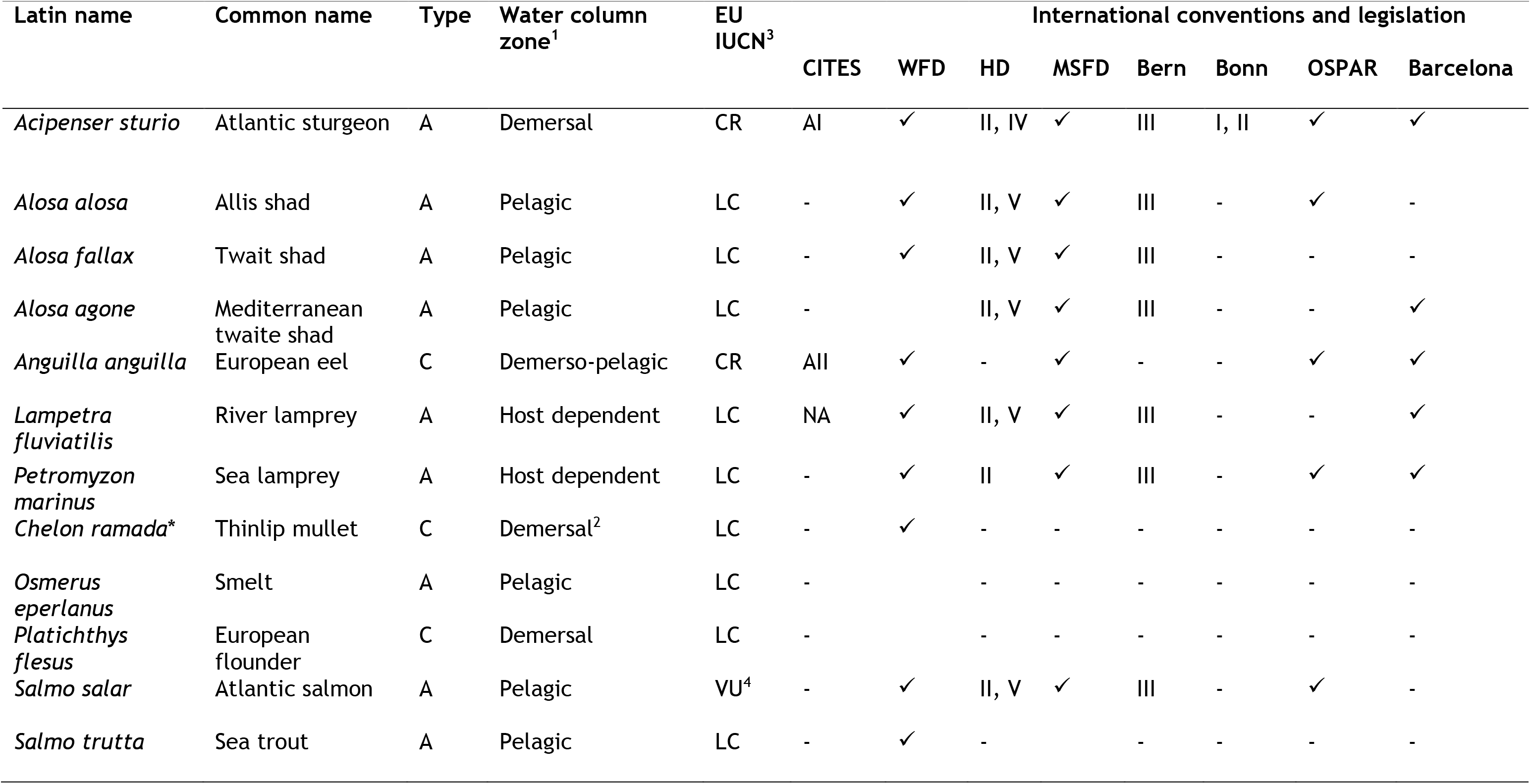
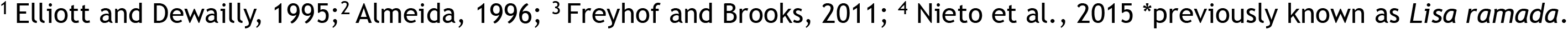
Diadromous fish observed within north-eastern Atlantic waters and their IUCN conservation status. A = Anadromous, C = Catadromous, - = not listed, CR = Critical, LC = Least Concern, VU = Vulnerable. Demersal species are found near the seabed, pelagic species occur in the open sea, demerso-pelagic species are species that migrate between these two water column zones, host dependent refers to lampreys that are parasitic species during their marine phase. ✓ = protected under the specific convention or Directive, letters specify the specific appendix the species is protected under.

Under the EU data collection framework, Member States are required to collect fisheries bycatch data through an onboard observer program (Cornou et al., 2015). Fisheries data is a rich set of information which provides year-round catch information and can be a valuable source of information for data-deficient species (e.g., Baum et al., 2003; Bisch et al., 2022; Elliott et al., 2020b). Bycatch data for protected species can be difficult to access, therefore fisheries observer data could help understand the distribution and bycatch mortality of data-poor species and meet EU Habitat Directive and MSFD (92/43/EEC; 2008/56/EC) requirements. Use of fisheries data does, however, require biases from the different gear types, the targeted nature of fishing, and un-balanced sampling to be taken into consideration (e.g. Alglave et al., 2022; Bourdaud et al., 2017).

Species Distribution Models (SDMs) provide a means to ‘fill in the gaps’ and provide complete coverage maps on which to base conservation and management decisions (Guisan and Thuiller, 2005; Guisan and Zimmermann, 2000; Leathwick et al., 2005). Modelling the distribution of diadromous fish could help improve protection measures for threatened and data-poor (insufficient biological information to determine the current exploitation status (Berkson and Thorson, 2015; Prince and Hordyk, 2019)) diadromous fish within existing MPAs. SDMs could also be used to meet Descriptor 1 (Biodiversity – habitat extent) requirements under the MSFD (2008/56/EC). However, building diadromous fish at sea distribution models remains challenging because most of these fish are either IUCN red listed at a national level (www.nationalredlist.org) or data-poor (Limburg and Waldman, 2009; Merg et al., 2020). Modelling the distribution of data-poor and rare species, can lead to underestimation of their true distribution due to imperfect detection (uncertainty in the presence of a species within a site where the data was recorded) (Guillera-Arroita, 2017; MacKenzie et al., 2002).

To model rare and data-poor species’ distribution, specific statistical methods are required to combine several sources of data that may not be initially designed to study the species distribution (Engler et al., 2004; Lomba et al., 2010; Simmonds et al., 2020). Using such data requires correction for detection bias arising from observer error, species rarity and variability in environmental conditions (Guillera-Arroita, 2017; Kellner and Swihart, 2014; MacKenzie et al., 2002) to help determine sites where the probability of presence of species is higher (Belmont et al., 2022). The low detectability of rare species can also result in a high proportion of false absences (Guillera-Arroita, 2017). Spatial and temporal variation in observation effort is another important source of heterogeneity in the observations (Guillera-Arroita, 2017; Kellner and Swihart, 2014; Meyer et al., 2011). Integrated hierarchical statistical model for species distributions have a great potential to combine multiple sources of data to infer a single latent field of abundance or presence/absence, and to enhance rare species distribution models for conservation and management purposes (Engler et al., 2004; Lomba et al., 2010; Simmonds et al., 2020). Another common problem which can arise when modelling species distribution is that data can be autocorrelated (more similar when closer together). Residual spatial autocorrelation can arise from population demographic processes or the influence of unobserved variables. Ignoring spatial autocorrelation can lead to inaccurate parameter estimations (Latimer et al., 2006). Developing models that explicitly account for spatial autocorrelation in the latent field of presence/absence is, therefore, necessary for accurate inferences on the distribution.

Given the lack of knowledge on the distribution of most diadromous fish during their marine life history stages, and the need to ensure they are sufficiently protected, our objective was to model the current distribution (2003-2019) of diadromous fish at sea. In addition, we wanted to quantify the risk of bycatch from different gear types and assess the value of Habitat Directive MPAs for their conservation. Since at sea targeted surveys do not exist for these species, we collated a database of 168 904 hauls from fisheries dependent (French fisheries bycatch observer data) and independent data (scientific survey data undertaken to assess fish populations) in eastern Atlantic and French Mediterranean waters. A Bayesian site-occupancy model, taking into account spatial autocorrelation in the probability of presence, and imperfect detection through gear type, was used for these data-poor species (Dormann et al., 2007; Latimer et al., 2006; MacKenzie et al., 2002; Moriarty et al., 2020). From the distribution models we wanted to answer the following questions: 1) Is the distribution of diadromous fish at sea connected to existing knowledge on their freshwater habitats? 2) Do different gears have different catchability, and does this provide information on the risk of bycatch of diadromous fishes? 3) Could existing Habitat Directive MPAs protect diadromous fish if specific measures are implemented?

## 2. Method

### 2.1. Combining fisheries independent and dependent surveys

Fisheries dependent and independent data from 1965 to 2019 were collated within eastern Atlantic waters (Greater North Sea, Celtic Sea, Bay of Biscay and the Iberian coast) and French Mediterranean waters). Scientific bottom trawl surveys were extracted from the International Council for the Exploration of the Sea (ICES) Database of Trawl Surveys (DATRAS) portal (https://www.ices.dk/data/data-portals/Pages/DATRAS.aspx; Refer to Elliott et al., 2021 Table S1; Data in brief Table S1 - S2) and French Metropolitan scientific surveys not submitted to ICES DATRAS were obtained from the Institut Français de Recherche pour l’Exploitation de la Mer (IFREMER) (https://campagnes.flotteoceanographique.fr/campaign). Fisheries-dependent data came from French fisheries observer data (ObsMer data; Cornou et al., 2015) that began in 2003. According to the sampling plan, fisheries observers sample fishing vessels and fishing operations when on board (Fauconnet et al., 2015). ObsMer data is held by IFREMER and available on request from the French Ministry of Fisheries and Aquaculture (Direction des Pêches Maritimes et de l’Aquaculture; Refer to Elliott et al, 2021 Table S1; Data in brief Table S1).

From the database, diadromous fish presence and absence, length, gear type, spatial location and year and month of capture were extracted (Fig. 1; Fig. S1). Due to missing data and insufficient information on the length of hauls, size of vessels, mesh size, etc., effort was not possible to calculate. Details of gear types and surveys that were used for the analysis can be found within the supplementary materials of Elliott et al., (2021).

**Fig. 1.**
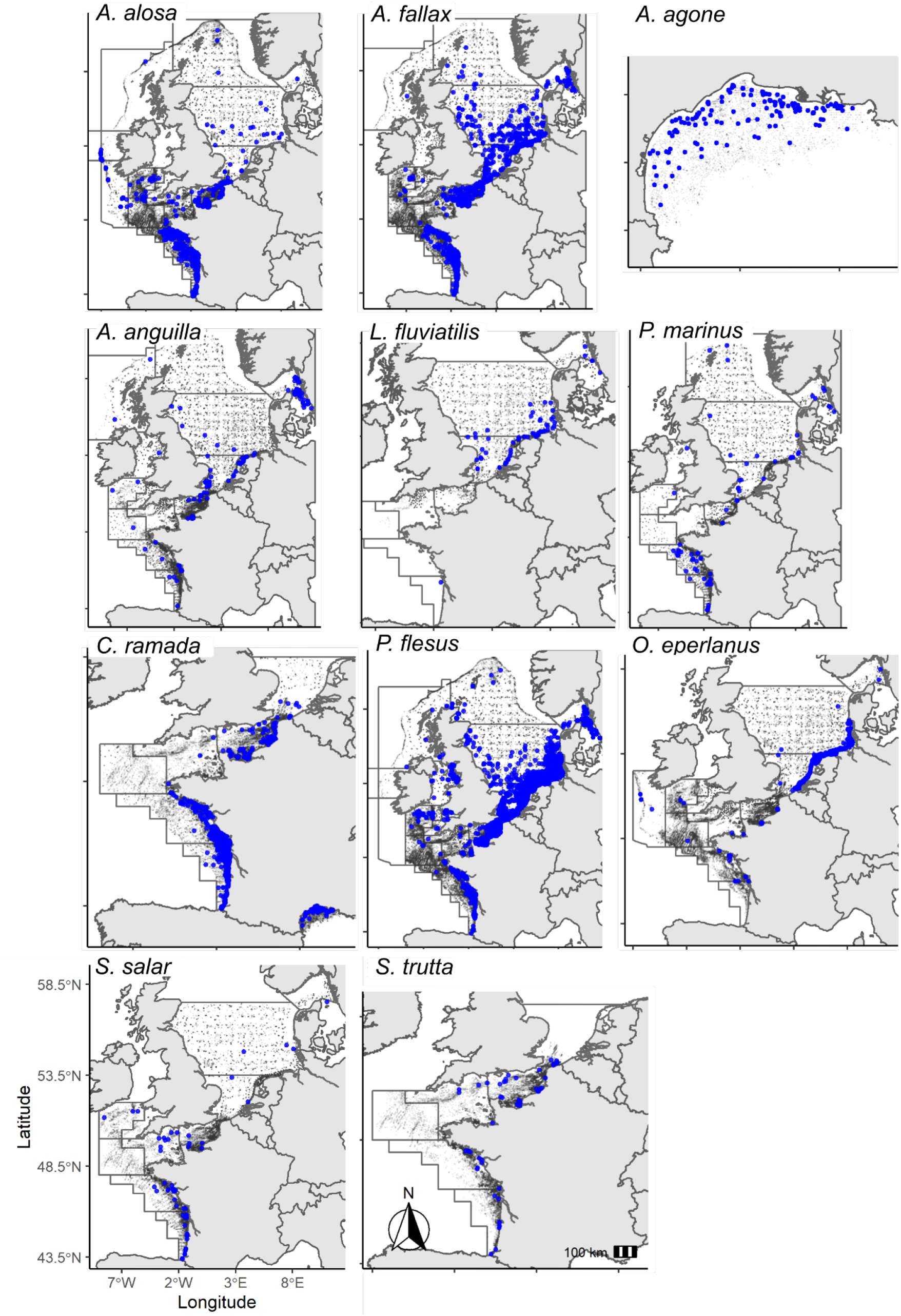
Diadromous fish presence (blue dots) and absence (light grey dots) data used to model their distribution (2003-2019). Dark grey lines indicate International Council for the Exploration of the Sea statistical divisions.

Prior to model analysis significant data cleaning was undertaken to standardise the surveys and extract relevant data to perform statistical analysis (e.g., converting gear types to gear categories, converting alike variables to the same units, etc.; see details within the Data in brief; Appendix A). Data from 2003 were used for spatial analysis, since a large proportion (74%) of the data came from the ObsMer dataset which started in 2003. In addition, prior to this data few presences were observed.

Since the proportion of zeros to presences was very uneven, which is not ideal for predictive modelling (Fielding and Bell, 1997), all surveys, gear types, target species (the intended catch for a particular fishery - from the ObsMer dataset) and ICES statistical divisions (Fig. 1) without diadromous fish presence, were deleted before modelling (Fig. S2 contains a spatial map of all hauls per gear type). Depths outside the ranges the species were observed were also removed. To avoid targeted fishing bias, all species that were targeted were removed. The latter included 2 European eels, *Anguilla anguilla*, 33 thinlip mullets, *Chelon ramada* and 41 flat fish which may have included European flounders, *Platichthys flesus* presences. All species were modelled other than the European sturgeon, *Acipenser sturio* because of too few presences (11 presences from 2003 – 2019). Note, for *L. fluviatilis*, one individual was caught by Set Longline within the Bay of Biscay. Due to its isolated nature of this capture, this caused problems when modelling (over detection of the line caught individual). Line gear type was therefore removed from the hierarchical modelling process.

### 2.2. Environmental predictor variables

Six environmental variables were considered as potential predictors of the presence of diadromous fish (seabed depth; distance from coast; sediment type; salinity; net primary production; and sea surface temperature; Table 2; Fig. S3). Depth and distance from the coast are thought to indicate key diadromous fish migratory periods (Taverny et al., 2012; Taverny and Elie, 2001; Trancart et al., 2014). Salinity and temperature are known to have direct physiological effects on diadromous fish, and their changes can indicate migration timing (Arevalo et al., 2020; Trancart et al., 2014). Net primary production is the gross primary production by autotrophs over the rate at which they respire, which is a measure of marine ecosystem functioning (ERSEM, 2020). Sediment type can be considered a proxy for food availability and shelter (Elliott et al., 2016; Trancart et al., 2014). As of result of too few presences and less information on length than the presence of individuals, distribution changes in length at stage, seasonal life history stage migration, and changes in habitat occupancy over the years were not considered.

**Table 2.**
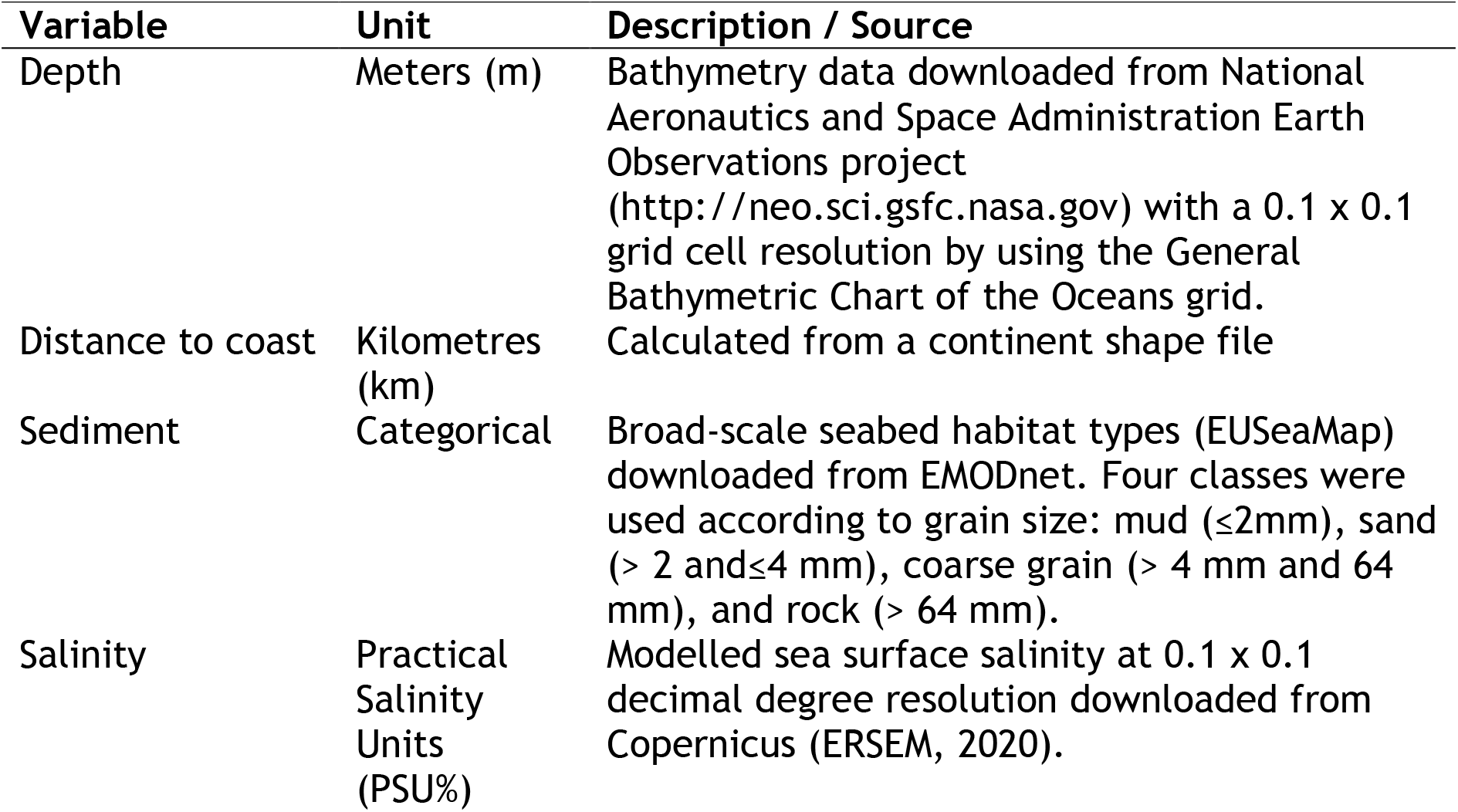
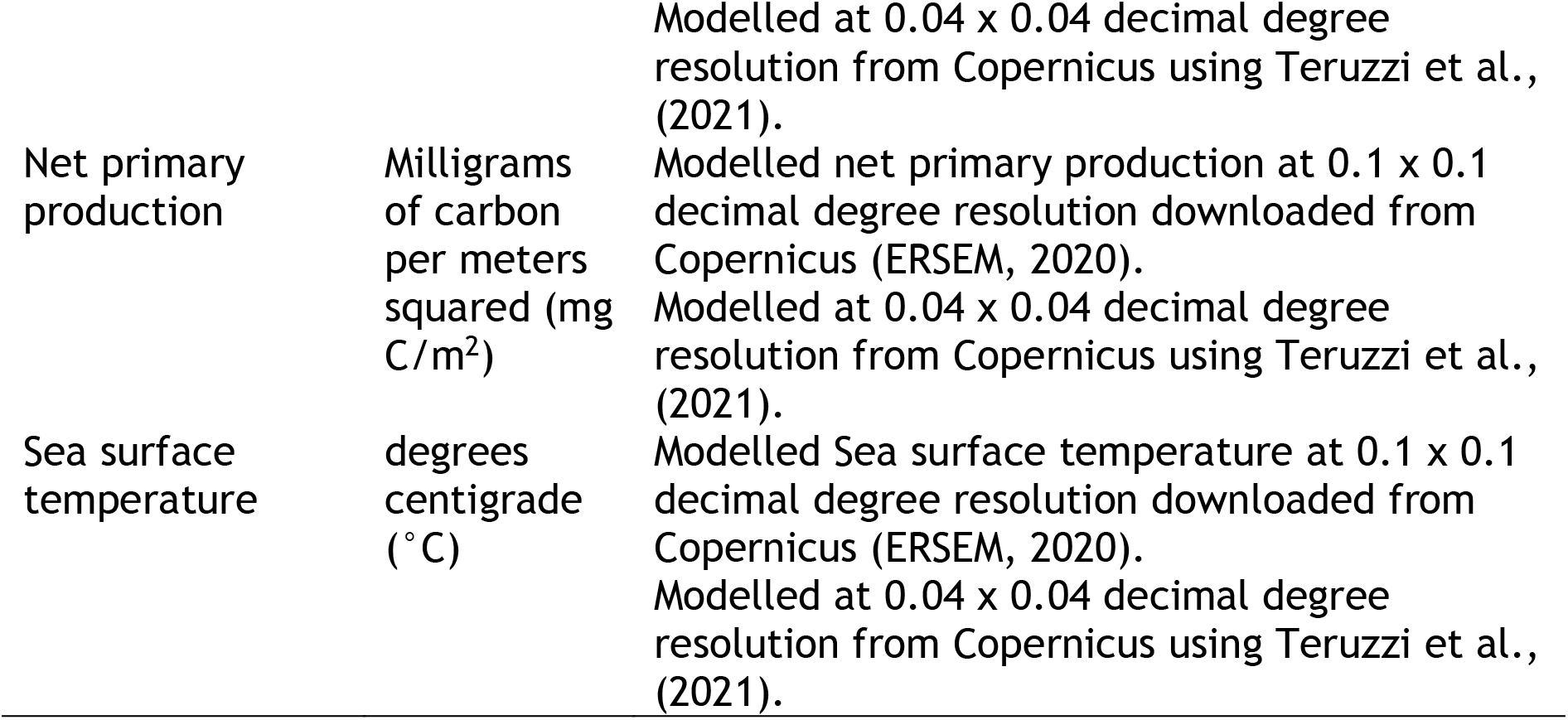
Environmental predictors used to analyse diadromous fish distribution. For all species except *Alosa agone*, variables were extracted at a 0.1 × 0.1 decimal degree resolution. *A. agone* environmental variables were extracted at a 0.04 decimal degree resolution.

To select the model of best fit, variables that were not collinear (Pearson’s correlation coefficient< 0.5 or >−0.5 and a Variance Inflation Factor < 2) (Dormann et al., 2013) were kept. Depth and distance from coast, and sea surface temperature and net primary production were collinear. Since most species occupied a wide latitudinal range and seasonal or yearly temperature effects were not modelled, net primary production was used instead of sea surface temperature. To identify the model of best fit a backwards stepwise model selection process was undertaken, ensuring distance from coast and depth were not in the same model. The model with the lowest deviance information criterion, which includes a penalty factor for the number of parameters, was selected (Spiegelhalter et al., 2002).

### 2.3. Statistical analysis

A Bayesian hierarchical site occupancy Gaussian intrinsic Conditional Autoregressive (iCAR) model was built independently for each species. Hierarchical models have the advantages of separating the ecological process (here, the habitat suitability model, that is the spatial distribution of the probability of presence/absence, including spatial autocorrelation) from the observation process (here, the imperfect detection and its variability among gears or other factors) (Dormann et al., 2007; Isaac et al., 2020; Latimer et al., 2006; MacKenzie et al., 2002). The fundamental concept of the model is that the true presence/absence spatial field (including spatial autocorrelation) is modelled on a grid, each grid cell being associated with multiple observation events with imperfect detection, with detectability that may vary depending upon the fishing gear.

The site occupancy model integrates two processes:

i. Latent ecological process (habitat suitability) The latent field of probability of presence is modelled on a grid as a function of environmental predictors and by explicitly accounting for spatial autocorrelation:

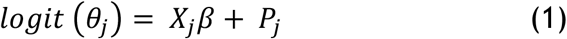

with *θ*_*j*_ the latent probability of presence (the habitat suitability) within grid cell cell *j*, modelled in the logit scale as a function of environmental predictors *X*_*j*_, with *β* a vector of fixed effect describing how much the environmental predictor contribute to the suitability process. The spatial autocorrelation *P*_*j*_ is modelled at the scale of the grid cell as an iCAR random effect (2):

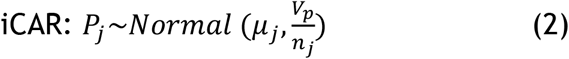

with *μ*_*j*_ the mean of the random effects calculated over the *n*_*j*_ cells considered in the neighbour of cell *j* (*n*_*j*_ is the number of the neighbours which is 8 in most cases, and <8 when the cell is on the boundary of the spatial domain), *V*_*p*_ the variance of the spatial random effect. A site is defined here as the area/volume covered during a given sampling/fishing operation. Any site *i* within the cell *j* as the same latent probability of presence *θ*_*j*_.
ii. Observation process (detection): Conditionally upon the latent probability of presence/absence in each grid cell *j* as describe above, the multiple detection events *y*_*j*,*i*_ (data 0/1) associated to the same grid cell *j* are modelled as mutually independent two steps Bernoulli process with detectability *δ*_*j*,*i*_:

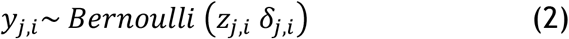

with

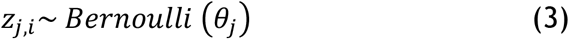

and

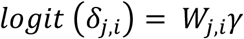

where *δ*_*j*,*i*_ denotes the probability of detecting the species at site *i* within cell *j*, modelled as a fixed effect of the gear type associated with observation *i*. *W*_*j*,*i*_ is the raw vector of the design matrix, and *γ* is the vector of the gear effects, so as *W*_*j*,*i*_*γ* is the gear affect associated with observation at site *i*.

All SDMs were undertaken using the ‘hSDM’ package (Vieilledent et al., 2014) using the ‘mod.hSDM.siteocc.iCAR’ function in R version 3.6.3 (R Code Team 2018). The prior distribution for the variance of the spatial random effect followed weakly informative uniform distribution. The effect of the detectability from gear and from the suitability process was modelled as fixed effects drawn from an weakly informative Gaussian prior centred at zero with a fixed standard deviation of 2 (Gelman et al., 2008; Northrup and Gerber, 2018). Each model was run with 50 000 Gibbs iterations, five Markov chain Monte Carlo simulations and a burn-in phase of 50 000 iterations.

### 2.4. Grid size and spatial distribution

A critical issue in our modelling framework is the choice of the grid size. Since the latent field of presence/absence is modelled at the scale of the grid cell (eq. 1), the larger the grid resolution the higher the probability to get at least one presence within a grid cell. A finer grid cell will limit the quantity of available observations falling within each cell, which therefore increases difficulty in model fitting. We selected the finest grid as possible that allow the model to converge. For almost all species modelled within Eastern Atlantic waters, a grid of 0.1×0.1 decimal degrees was used. Because of the scarcity of data for Lampreys, the model did not converge with such a resolution and a coarser resolution of 0.2×0.2 decimal degrees was therefore used for these species. For *A. agone*, that only occurs within the Mediterranean, a grid resolution of 0.04 × 0.04 decimal degrees was used because of the smaller area encompassed. This spatial resolution (0.04 × 0.04 decimal degrees) was not used for other species since it was not available at the scale required within Eastern Atlantic waters. All environmental variables were then aggregated and modelled to their relevant spatial resolution. There were insufficient presences within the Mediterranean for *A. anguilla*, *P. marinus*, *C. ramada*, *P. flesus* predictions (issues with model convergence). Predictions within the Mediterranean for these were therefore excluded (Fig. 2).

**Fig. 2.**
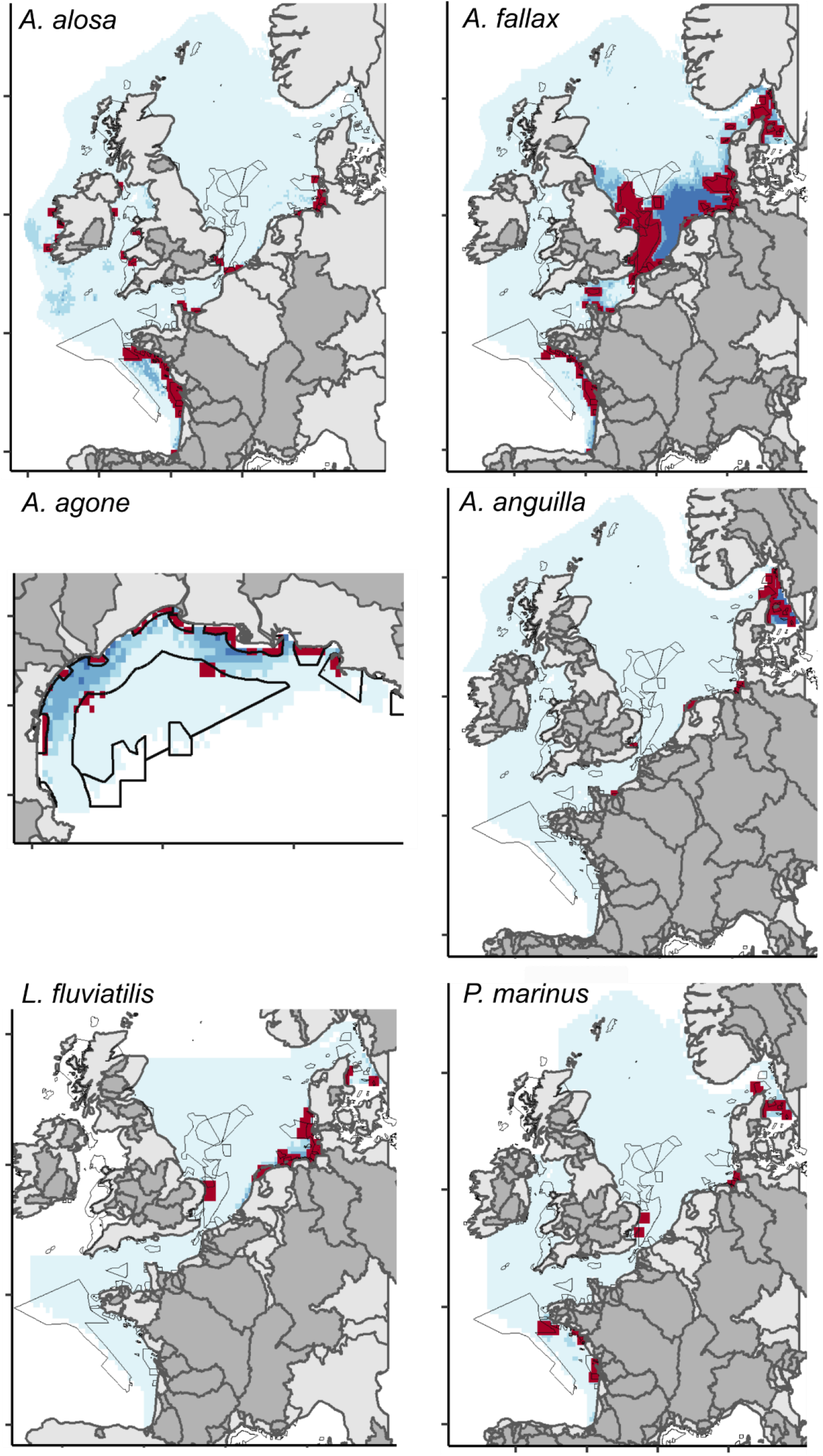
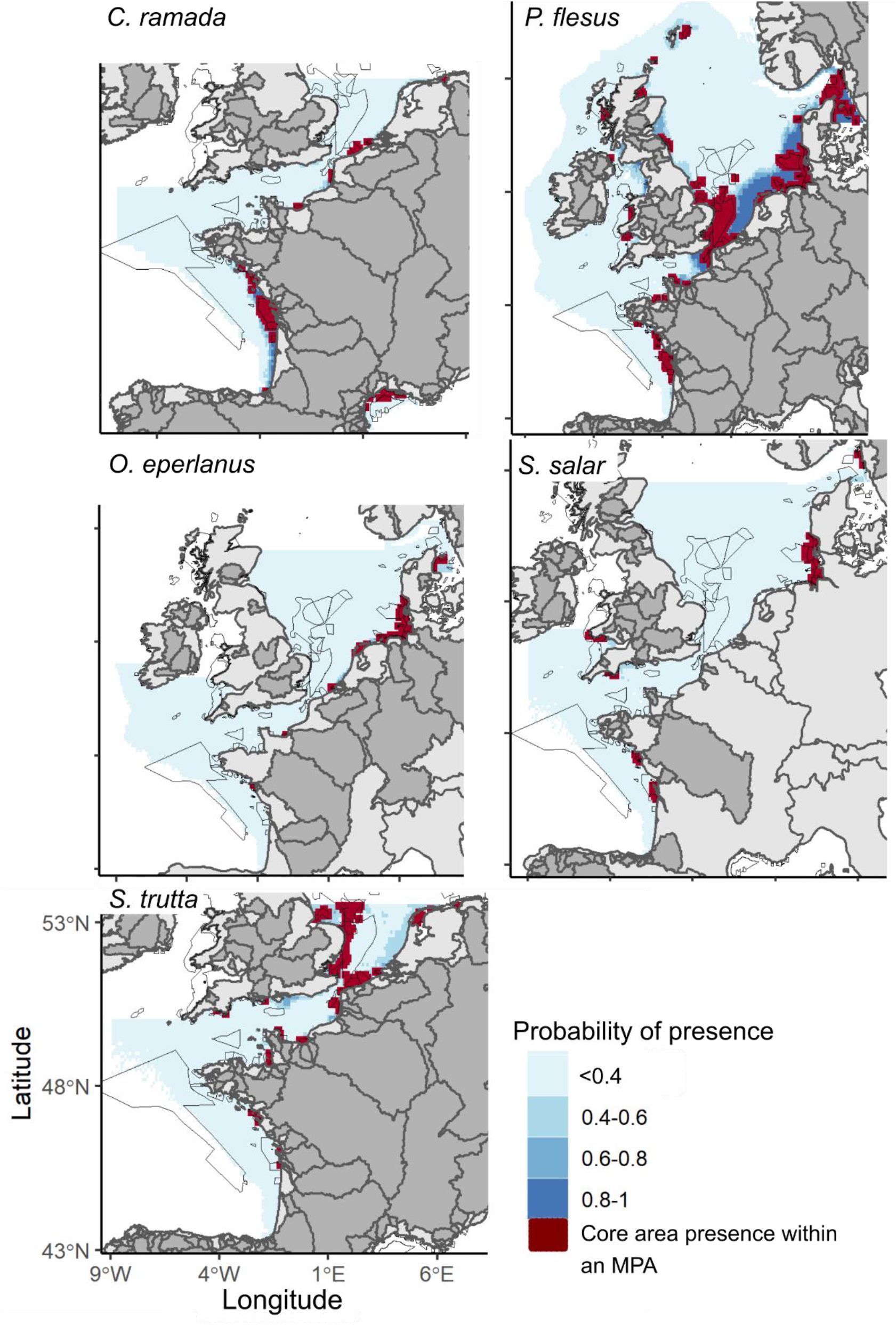
Diadromous fish hierarchical species distribution model probability of presence. Darker blue shades = SDM probability of presence > 0.4, light blue shade = SDM probability of presence ≤ 0.4. Habitat Directive Marine protected Areas (MPAs) outlined in black. MPAs of relevance to core (probability of presence > 0.4) diadromous fish distribution, highlighted in dark red. Dark red cells are larger than the gridded cells to ensure smaller core areas within MPAs are visible. Darker grey terrestrial areas represent river basins which the diadromous fish migratory populations have been observed (EuroDiad V.4). Note, no data were available within the EuroDiad database from Dutch river basins.

### 2.5. Model evaluation

The probability of presence was calculated from the mean of the posterior distribution of the latent field *θ*_*j*_. A threshold of 0.4 was used to bin estimated probability of presence into binary presence/absence values. A threshold of 0.4 was used as a compromise given the rarity of the species and accuracy following examining various thresholds and plotting positive predicted values. The model performance was then evaluated using sensitivity (proportion of correctly classified presences), specificity (proportion of correctly classified absences) and percent correct classification (proportion of presences and absences correctly using the R package ‘PresenceAbsence’ (Freeman and Moisen, 2008a; Fielding and Bell, 1997; Table 3). Max sensitivity specificity which is often used for rare species distribution models (Freeman and Moisen, 2008b; Jiménez-Valverde and Lobo, 2007), was not used since false presences was very high when applying this metric. Additionally, our SDM objective was to try and minimise false presences and absences within the distribution models. The threshold independent Area under the receiver operating curve was also used to evaluate the model performance. These four metrics were used to better understand the accuracy of the model since, some metrics can be biased with low prevalence data and where imperfect detection can arise (Allouche et al., 2006; Fourcade et al., 2018; Guillera-Arroita, 2017; Leroy et al., 2018).

**Table 3.** Hierarchical species distribution model outputs. All models contained gear as part of the observation process. AUC = Area Under the receiver operating Curve score, PCC = Percent Correct Classification, NetPP = Net Primary Production. Max depth is the maximum depth at which the dataset was cut. Note most environmental variables (all except distance and depth) were not available within very coastal areas (e.g., within estuaries and inlets). Numerous presences and absences were therefore lost when extracting the environmental variables for modelling.

| Species | Model selected | Hauls | Presence | Absence (%) | Max depth (m) | Sensitivity | Specificity | AUC | PCC | Cell size (decimal degrees) |
| --- | --- | --- | --- | --- | --- | --- | --- | --- | --- | --- |
| *A. alosa* | Depth + Salinity + NetPP + Sediment | 86 292 | 802 | 99 | 300 | 0.79 | 0.92 | 0.95 | 0.95 | 0.1x0.1 |
| *A. fallax* | Distance + Salinity + NetPP + Sediment | 91 135 | 1385 | 98 | 300 | 0.88 | 0.81 | 0.90 | 0.82 | 0.1x0.1 |
| *A. agone* | Depth + Salinity + Sediment | 5086 | 176 | 97 | 200 | 0.81 | 0.74 | 0.86 | 0.75 | 0.04x0.04 |
| *A. anguilla* | Depth + Salinity + NetPP | 54 124 | 176 | 99 | 250 | 0.53 | 0.99 | 0.98 | 0.98 | 0.1x0.1 |
| *L. fluviatilis* | Distance + Salinity | 16 536 | 68 | 99 | 150 | 0.71 | 0.95 | 0.96 | 0.94 | 0.2x0.2 |
| *P. marinus* | Distance + Salinity | 28 582 | 74 | 99 | 300 | 0.60 | 0.98 | 0.94 | 0.97 | 0.2x0.2 |
| *C. ramada* | Distance + Salinity + Netpp + Sediment | 53 083 | 925 | 98 | 300 | 0.84 | 0.93 | 0.96 | 0.92 | 0.1x0.1 |
| *P. flesus* | Distance + Salinity + NetPP + Sediment | 92 400 | 5394 | 94 | 300 | 0.90 | 0.93 | 0.97 | 0.93 | 0.1x0.1 |
| *O. eperlanus* | Depth + Salinity + NetPP | 48 877 | 1035 | 98 | 450 | 0.81 | 0.99 | 0.99 | 0.99 | 0.1x0.1 |
| *S. salar* | Depth + Salinity + NetPP | 36 008 | 68 | 99 | 150 | 0.66 | 0.98 | 0.95 | 0.98 | 0.1x0.1 |
| *S. trutta* | Depth + Salinity + NetPP | 31 754 | 63 | 99 | 150 | 0.71 | 0.94 | 0.93 | 0.93 | 0.1x0.1 |

So that areas with probabilities of presence above the threshold could be easily identified for management purposes, all predictions below 0.4 were mapped with a uniform light blue colour (Fig. 2). Uncertainty maps for each model were also produced to gage a better idea of areas of prediction with higher uncertainty (Loiselle et al., 2003; Spiegelhalter et al., 2002). The uncertainty maps were calculated by the mean coefficient of variation from the posterior distribution of spatial parameters in the model of best fit for each species (Lambert et al., 2020; Latimer et al., 2006). Probabilities of prediction and uncertainty above 0.4 were classified into 0.2 units to highlight areas of higher and lower probability of presence more clearly (Fig. 2; Fig. S4). As a result of the low diadromous fish presence, splitting the data into learning/validation samples led to insufficient presence in the learning data set. Cross validation was therefore not undertaken.

### 2.6. Diadromous fish river basin presence

The predicted distributions were visually compared to the distribution of diadromous fish within their freshwater habitats using the EuroDiad database (version 4; https://data.inrae.fr/dataset.xhtml?persistentId=doi:10.15454/IVVAIC; Barber-O’Malley et al., 2022). From EuroDiad database, migrating diadromous fish population presence within the study areas our model encompassed, were extracted using ‘recent’ literature citing from 1951 to 2020 (see Barber-O’Malley et al., (2022) and Béguer et al., (2007) for more details on the database) and mapped with the hierarchical SDM results (Fig. 2). The EuroDiad4 database was used as opposed to IUCN red list map distribution since it has been peer reviewed and it is more up to date.

### 2.7. Habitat Directive MPAs of relevance to diadromous fish

‘Core areas’ with a probability of presence >0.4 were selected and intersected with European Union Natura 2000 Habitat Directive Sites of Community Importance (https://www.eea.europa.eu/data-and-maps/data/natura-12). A threshold of > 0.4 probability of presence was selected as a result of the low probability of presence of diadromous fish (particularly Habitat Directive protected species), and to ensure consistency with the model evaluation threshold. Since only one Site of Community Importance was submitted to the European Union from the United Kingdom, Special Areas of Conservation were taken into consideration (https://jncc.gov.uk/our-work/special-areas-of-conservation-overview/). For each species, the number of MPAs (Sites of Community Importance and Special Areas of Conservation) and the proportion of MPAs of relevance to diadromous fish (core areas within an MPA) were recorded. The extent of the core area within the MPAs was not calculated as this would be dependent of the model grid used (Table 3). Furthermore, numerous MPAs were smaller than the grid.

## 3. Results

### 3.1. Diadromous fish hierarchical SDM

For all species the Area under the receiver operating curve and Percent Correct Classification performed very well (≥75%; Table 3). Correct presence classification from the confusion matrices was consistently lower (<15%) than that of correct absence classification (≥64%), and false absences were lower (≤3%) than false presences (≤22%) because of presence absence imbalance (Table 3; Table S1). The later led to on average better specificity than sensitivity scores (Table 3).

*P. flesus* was the most prevalent species modelled (5394 presences), whereas salmonids the least (*S. salar* = 68 presences and *S. trutta* 63 presences) (Table 3). SDMs that had lower presences (salmonids and lampreys) did not predict as well (lower sensitivity and higher uncertainty maps) as species with higher presence (shads, *C. ramada*, *P. flesus* and *O. eperlanus*). In addition, species that had a more disperse distribution (e.g., *A. anguilla* and *S. salar*) did not predict as well (sensitivity ≤ 0.66 for both species) as those with a more aggregated distribution (*O. eperlanus* sensitivity = 0.81) (Table 3; Fig. 1 and 2). Uncertainty maps showed that in general, areas of high and low predicted probability of presence had lower uncertainty (Fig. S4).

All species were observed to have a relatively shallow (<300m depth) coastal distribution (~<300 km from the coast; Fig. 2 and 3). The main predictor variables for diadromous fish habitats were depth or distance from coast and salinity (Fig. 3). Since sufficient *C. ramada* were observed within eastern Atlantic and Mediterranean waters to model, two salinity peaks are observed corresponding to the different salinity ranges of the two regions (Fig. 3).

**Fig. 3.**
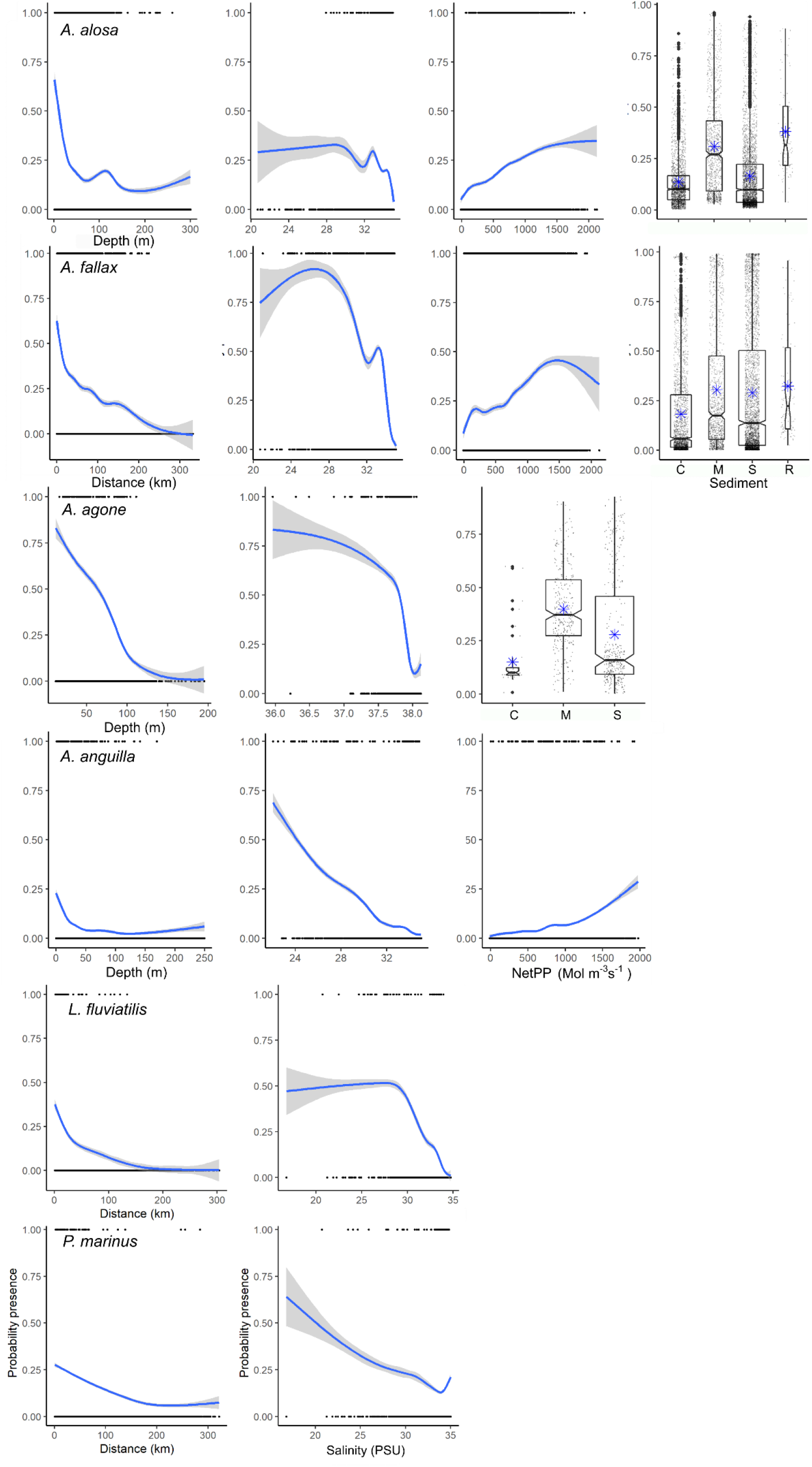
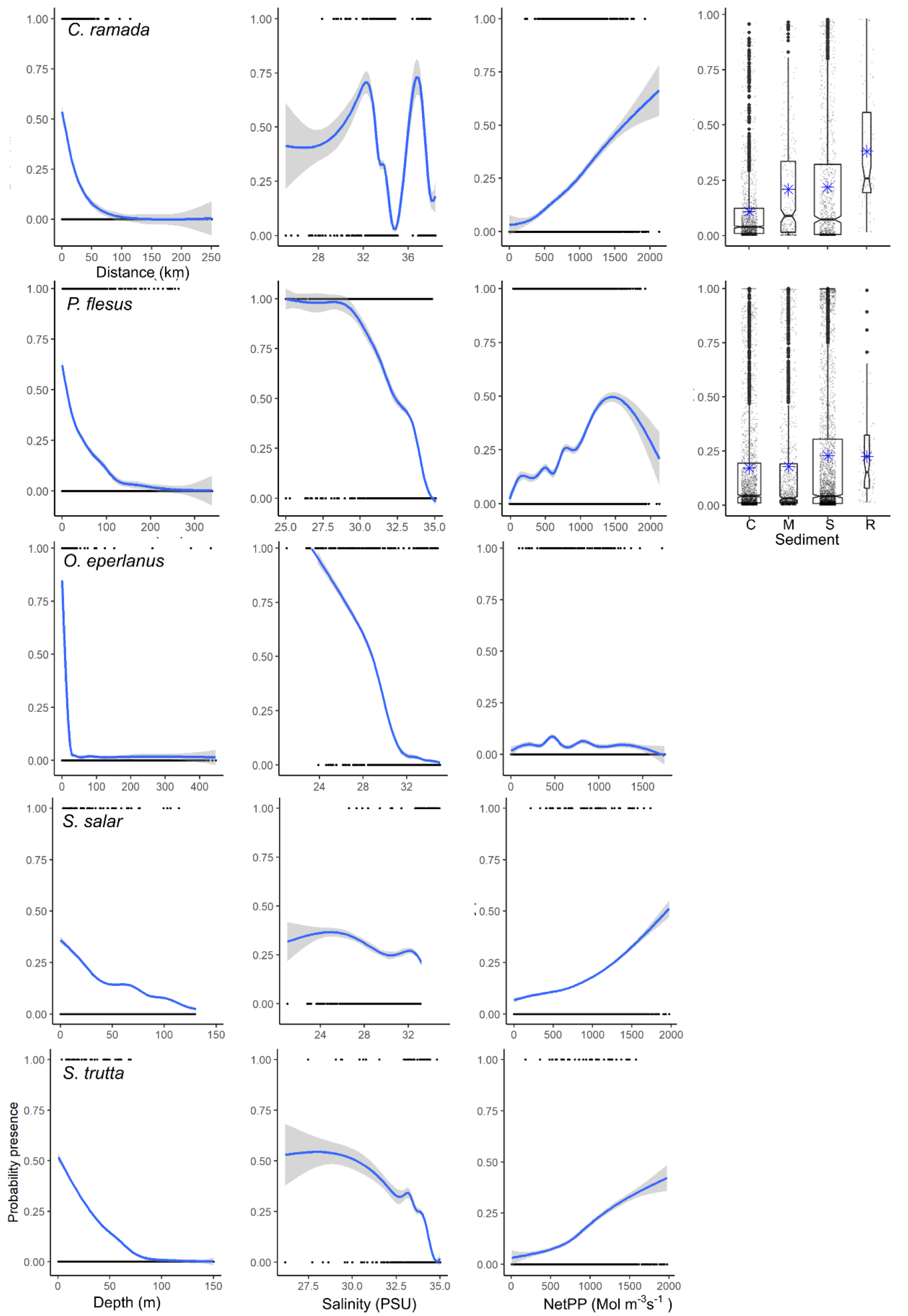
Diadromous fish hierarchical species distribution model environmental predictor effects. NetPP = Net Primary Production, C = Course grain, M = Mud, S = Sand, R = Rock. The solid line represents the smooth function estimate; the shaded region represents the approximate 95% credibility interval. Boxplot blue stars represent means and small black dots the predictions.

### 3.2. Diadromous fish river basin connectivity

2717 references contained within the EuroDiad database were used to map the 11 diadromous fish presence within the river basin. Core distributions of diadromous fish at sea largely matched with the freshwater areas of presence (Fig. 2). The spatial extent of diadromous fish at sea was often less than that observed within river basins (e.g., lampreys, *C. ramada*, *O. eperlanus* and *S. trutta*; Fig. 2). It should be noted that there are gaps in the EuroDiad database (refer to Barber-O’Malley et al., (2022)). For example, no diadromous fish presence data were available within Dutch freshwater habitats, and salmon populations are known to exist within Danish waters (de Groot et al., 2012; Maes et al., 2007; Rikardsen et al., 2021). Presences of *A. alosa* and *A. fallax* within the Mediterranean from the EuroDiad database will have been due to historic classification (Bagliniere and Elie, 2000; Keith et al., 2020).

### 3.3. Gear type detectability

All species were caught by a range of gear categories ranging from benthic, demersal and pelagic mobile trawls, seine nets, line gear types (Fig. 4; Table S2). Some general patterns were, however, evident. Line gear types caught the least diadromous fish (2% (weighted), 17 presences over ~2887 hauls), whereas demersal mobile trawl gear types (otter beam trawl and otter twin trawl) caught the most (67%, 7409 presences over ~82 531 hauls; Fig. 4; Table S2). Shad had a higher detectability with static and demersal mobile gears. *A. anguilla* were detected by a wide range of gear types. Lamprey were mainly detected by mobile trawl gears. *P. flesus* and *O. eperlanus* had a higher detectability with demersal and benthic mobile gear, and salmonid had a higher detectability with static nets and pelagic gears. Rarer species and species that were caught by a wider range of gear types had lower detectability than more abundance species caught by fewer gear types (Fig. 4). On average static nets (e.g., trammel net and set gillnets) and pelagic mobile gears (e.g., Otter midwater trawls) caught larger diadromous fish than mobile demersal and benthic gear types (Fig. S1). Although the modelled species were not target species of the fisheries within the observer database, some species were landed (Table 4).

**Fig. 4.**
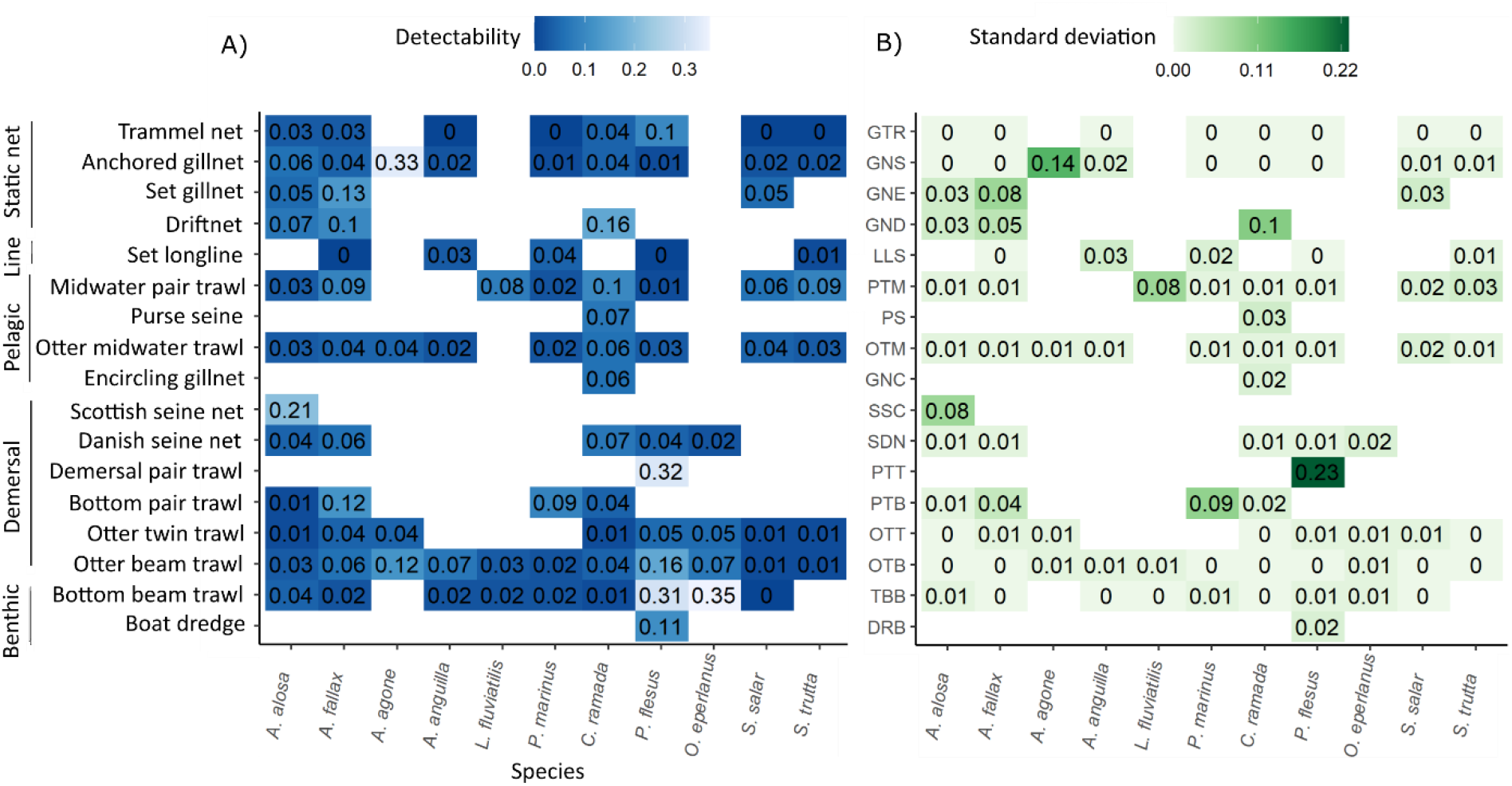
Matrix of the posterior mean (marginal posterior distribution) for gear A) detectability (inverse logit of; the probability to detect the presence of fish when they are present) and B) its standard deviation. Lighter blue shades represent higher detectability (A) and darker green shades higher detectability standard deviation (B).

**Table 4.** Landed and discarded diadromous fish from the ObsMer data. Excluding targeted catches. Proportion landed = total landed / landed + discards.

| Species | Landed | Discard | Hauls | Proportion landed |
| --- | --- | --- | --- | --- |
| *A. alosa* | 182 | 515 | 70 378 | 0.26 |
| *A. fallax* | 152 | 608 | 64 300 | 0.20 |
| *A. agone* | 28 | 69 | 7373 | 0.29 |
| *A. anguilla* | 10 | 8 | 28 194 | 0.56 |
| *L. fluviatilis* | 0 | 2 | 828 | 0.00 |
| *P. marinus* | 2 | 18 | 10 030 | 0.10 |
| *C. ramada* | 643 | 101 | 45 699 | 0.86 |
| *P. flesus* | 856 | 648 | 66 196 | 0.57 |
| *O. eperlanus* | 6 | 45 | 31 311 | 0.12 |
| *S. salar* | 29 | 27 | 26 009 | 0.52 |
| *S. trutta* | 43 | 18 | 31 666 | 0.70 |

### 3.4. Habitat Directive MPAs relevance

A total of 482 Habitat Directive MPAs have been designated within the area in which the species were modelled (Fig. 2). Despite the small area of core habitats relative to the model prediction area, for most Habitat Directive listed species, numerous MPAs overlap with species core habitats, indicating their potential for the protection of diadromous fish at sea (Table 5; Fig. 2).

**Table 5.**
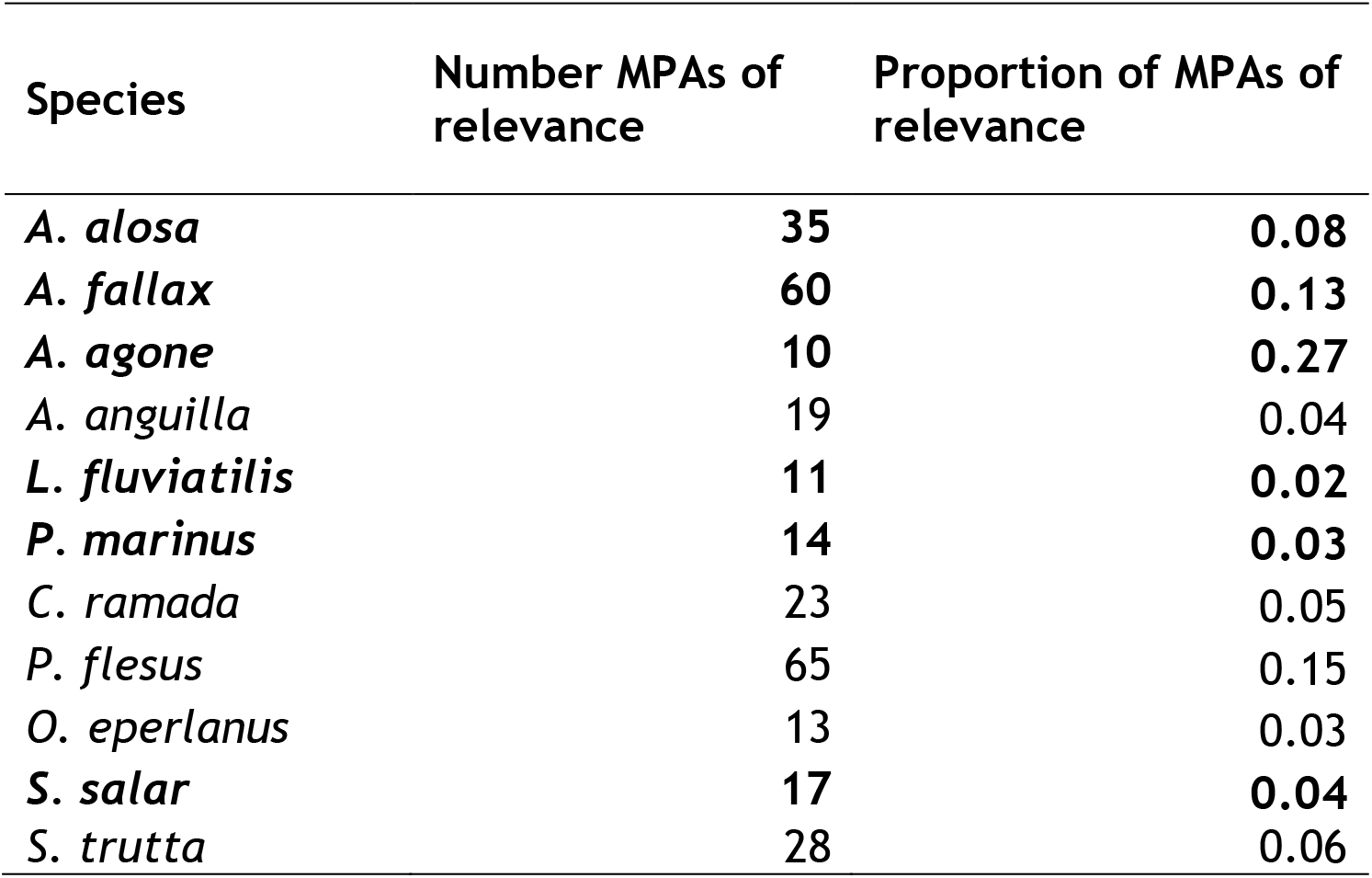
Habitat Directive Marine Protected Areas of relevance to core diadromous fish distribution (probability of presence >0.4). Diadromous fish listed under the Habitat Directive in bold.

## 4. Discussion

To our knowledge this is the first comprehensive study to model the distribution of diadromous fish at sea across eastern Atlantic and French Mediterranean waters. Using a hierarchical model allowed us to integrate an extensive database that we collated, containing both fisheries dependent and independent data to model the distribution of eleven rare diadromous fish over a wide spatial area. The model also enabled us to evaluate the probability to detect the presence of fish by the different gear types, and thus provide key information to evaluate the risk of bycatch of diadromous fish by commercial fisheries at sea.

Diadromous fish are bound to coastal areas because of their migratory behaviour. They are known to be fragile species because of the physiological and osmoregulatory changes required to undertake migrations between fresh water and marine environments (McDowall, 2009). Yet, coastal ecosystems, particularly within eastern Atlantic and Mediterranean waters, are subject to some of the highest anthropogenic impacts (Halpern et al., 2015; Korpinen et al., 2021; Lotze et al., 2006). Even though more data were collected within coastal areas, and the species are known to migrate from freshwater habitats, our study confirms that the investigated species were found to mainly concentrate in coastal and relatively shallow waters. The strong dependency of diadromous fish species to coastal areas, as evidenced in this study, reinforces the importance of investigating the vulnerability of diadromous fish to coastal anthropogenic pressures (Halpern et al., 2015; Limburg and Waldman, 2009; Lotze et al., 2006; Worm et al., 2006).

### 4.1. Hierarchical Bayesian SDM

On average, the eleven SDMs performed well. All the models had lower sensitivity and positive prediction values scores, because of low presence. Areas of both high probability and very low probability of presence had lower uncertainty, giving a more accurate picture of confidence in the distributions (e.g., higher certainty that most species are absent within waters further from the coast). In addition to problems with very low occurrence, the areas of higher uncertainty for certain species (e.g., *S. salar* and *S. trutta*, *A. alosa* and *A. fallax*), may be partially because of misidentification between these groups of diadromous fish. The core areas of probability of presence, in combination with the confidence maps provide a better understanding of confidence of these predictions. To improve problems of false presence predictions which might occur with species that are difficult to identify, Bayesian models incorporating false positive errors could be undertaken (e.g. Diana et al., 2021; Guillera-Arroita, 2017; Royle and Link, 2006).

Separating out the spatial random field of presence absence from the observations enabled us to explicitly consider the imperfect detection of the observation process. Since our database contained non-diadromous fish specific survey and catch data which sampled the same grid cell numerous times, the use of the site occupancy model enabled imperfect detection parameters to be estimated. Disregarding false absences would have led to biased inferences and over-stated parameter precision (Guillera-Arroita, 2017). The spatial random field of the probability of presence explicitly integrates spatial autocorrelation in the probability of presence, which limits the bias in inferences due to the non-random spatial repartition of observations.

Since there were few presences for most species, our results were sensitive to the choice of prior on the variance of the spatial random effect. We tested both the inverse-gamma or a uniform distribution (the two prior forms that are available using the hSDM package; Vieilledent et al., 2014) and found that the uniform distribution provided better result. Some alternative prior choice such as the half-Cauchy are advocated in the literature (Gelman, 2006), and future work should explore if other prior choice could provide more accurate inferences. Equally, for increased confidence in predictions of rare species, several different distribution models could be compared or an ensemble model (that uses the means of several models) implemented to stabilise inferences (Araújo and New, 2007; Latimer et al., 2006; Loiselle et al., 2003).

For conservation purposes low thresholds are advised for low occurring species such as diadromous fish (Freeman and Moisen, 2008a;b; Jiménez-Valverde and Lobo, 2007). However, when lowering the threshold an increase in false positives can occur (Freeman and Moisen, 2008b). Our objective for the distribution models was to minimise false presence and absences where possible, to provide more accurate predictions whilst trying to meet conservations needs and minimise potential impacts on sea-users any spatial protection measure that might be implemented (Domisch et al., 2018; Loiselle et al., 2003; Maxwell and Jennings, 2005). Lack of understanding in species distribution can lead to inefficient spatial protection measures (Wauchope et al., 2022) and resource conflict (Probst et al., 2021).

### 4.2. Diadromous fish river basin connectivity

The EuroDiad database is a large database that incorporates information on diadromous fish presence and abundance during their freshwater stage (Barber-O’Malley et al., 2022; Béguer et al., 2007). Using the Eurodiad database to provide large scale understanding of river basin presence of diadromous fish, we provide a first attempt at qualitatively linking diadromous fish freshwater and marine distributions within north-eastern Atlantic waters. Although the probabilities of presence for most species were low, and gaps within the EuroDiad database evident (e.g., gaps in diadromous fish presence from Denmark and Netherlands), connectivity with their freshwater habitats appears to be visible, indicating migration pathways.

It is thought *A. alosa* no longer exists within the Greater North Sea (Baglinière et al., 2003; Wilson and Veneranta, 2019). Presences observed may have been from misidentified *A. fallax*. The high probability of presence of *A. fallax* along the east coast of the UK may be because of suitable conditions for this species as few spawning rivers have been recorded along the east coast (Aprahamian et al., 1998). For *A. agone* probabilities of presence clearly match that of their watersheds. Predicted *A. anguilla* presences closely match tagged river presences from Righton et al., (2016) despite the gaps found in the EuroDiad database. Predicted presence of *L. fluviatilis* was higher along the north-west coast of Germany and Holland. Presences have been observed here, albeit in low numbers (Admiraal et al., 1993; Pavlov et al., 2017; Thiel and Salewski, 2003). *P. marinus* predicted presence was slightly more scattered which fits with its wide distribution including into deeper waters (Elliott et al., 2021; Lança et al., 2014). The very low occurrences of *A. anguilla*, and the lamprey species is also likely to be due to inadequate sampling methods used to study these species.

Probabilities of presence of *C. ramada*, *P. flesus* and *O. eperlanus* matched relatively well their known presence from the EuroDiad database. From the EuroDiad database, there are few *S. salar* river basin presences within the North Sea. However, *S. salar* are known to occur within the North Sea and migrate Northwards to their feeding grounds (Mork et al., 2012; Rikardsen et al., 2021). Occurrences observed were most likely 1 sea winter and 2 sea winters individuals returning to their natal rivers given their size ranges. From the rivers that are monitored, *S. trutta* distribution matched well their freshwater river occupancy (ICES, 2020). The few salmonids observed may be as a result of the relatively few pelagic trawls, in addition to misreporting (ICES, 2005).

Given the coastal habitat occupancy for most of these diadromous fish and declines observed (Limburg and Waldman, 2009; Merg et al., 2020), detailed analysis of the connectivity between both habitats is essential (Flitcroft et al., 2019; Lin et al., 2017; McDowall, 2009). Unfortunately, such a model was not possible here, because it would require more detailed knowledge on their freshwater habitat occupancy (i.e., numerous outlets diadromous fish were observed within proximity to, were not contained within the EuroDiad database) and their river fidelity. Furthermore, seasonal and stage specific SDMs, which would provide more information on their ontogenetic migration movements, were not possible because of the low prevalence of the data and dispersion. Developing future research to model the connectivity between both habitats is key to improving our understanding of the pressures faced during their migrations (Flitcroft et al., 2019; Lin et al., 2017; McDowall, 2009).

### 4.3. Gear type detectability

Even though little bycatch was observed, given a number of the diadromous fish studied here are threatened, even a small amount of bycatch may impact their populations (Dulvy et al., 2003). Furthermore, misreporting and illegal fishing is likely to remain (Elliott et al., 2020a; ICES, 2005; Stratoudakis et al., 2020; Worm et al., 2013). Bycatch detectability results from our models could help provide management advice so that gear types and areas with higher bycatch rate be avoided. Such advice can provide direct information for the MSFD and the Habitat Directive that requires the identification of gear types which may threaten protected species (1992/43/EEC; 2008/56/EC).

All species were caught by a wide range of gear types (e.g., benthic, demersal and pelagic trawls, and seine nets), despite the different gear categories being deployed at different depths. The species were probably caught by a range of gear categories because of their diel vertical migratory behaviour whilst at sea (Kristensen et al., 2018; Lança et al., 2014; Righton et al., 2016). In addition, certain gear types (e.g., trawls and static nets), can catch fish within both the demersal and pelagic water zones (He et al., 2021; Borges et al., 2008). Higher detectability from certain gear types were, however, observed. For example, shad were mainly caught by demersal mobile gear and static gear, salmonids static and pelagic mobile gear types, and *P. flesus* and *O.eperlanus* were largely captured by demersal and benthic mobile gear. These results broadly match the water column habitat occupancy these species are known to occupy and existing literature (ICES, 2005; Wilson and Veneranta, 2019). As it is thought that lampreys detach from their host upon capture (Elliott et al., 2021; Halliday, 1991), bycatch is likely to be minimal for these species.

To improve understanding of bycatch risk on a spatial level, fishing intensity by gear type should be mapped, ideally using Vessel Monitoring System data and overlaying it with the predicted distributions (Elliott et al., 2018; Quemmerais-Amice et al., 2020). Unfortunately, this information was not available to us. Additionally, since the fisheries observer data we had access to is only a small sample of the existing French fishing activity (Cornou et al., 2015), and it does not include other countries fishing effort, results would be biased.

### 4.4. MPA relevance

To improve protection of diadromous fish at sea, numerous mechanisms could be implemented. For example, identifying areas of high bycatch risk with measures to limit bycatch within these areas, improving diadromous fish migratory pathways, and spatial protection measures (Verhelst et al., 2021; Wilson and Veneranta, 2019). Few MPAs within European waters protect marine fish and instead focus on protecting seabed features with a few exceptions (e.g., protected seabirds and marine mammals). Therefore the only protection measure that exist for vulnerable marine fish is through the removal of targeted fishing (Dureuil et al., 2018; Probst et al., 2021; Stratoudakis et al., 2016). There is, however, increasing evidence that MPAs may have positive impacts on fish populations if appropriate management measures are implemented (Davies et al., 2021; Moland et al., 2013; Probst et al., 2021; Worm et al., 2006). The value of protected areas for highly migratory species has been questioned, but there is evidence that MPAs may have positive impacts on migratory species (Pendoley et al., 2014; Takashina and Mougi, 2014).

Our results show relatively high core area presence of protected diadromous fish within Habitat Directive MPAs. Most marine Sites of Community Importance have not been designated to protect Habitat Directive listed diadromous fish. Nonetheless, our results highlight the value of these MPAs for diadromous fish protection. Specific management measures limiting threats (e.g., limiting access to gear types with higher probability of capture) to diadromous fish within these areas, would be of benefit to their protection (Domisch et al., 2019; Stratoudakis et al., 2016). Detailed analysis of Sites of Community Importance and other categories of designated MPAs should be undertaken to evaluate their potential to protect diadromous fish.

### 4.5. Conclusion

Understanding the spatial distribution of species and their habitats is essential for effective management and conservation (Halpern et al., 2015; Worm et al., 2009, 2005). Much research has been carried out on the conservation of freshwater diadromous fish (Drouineau et al., 2018; Merg et al., 2020; Verhelst et al., 2021). Very little research has been dedicated to the value of marine and estuary habitats for the conservation of these species and their risk of bycatch (Feunteun, 2002; Flitcroft et al., 2019; Lin et al., 2017). Here we provide an insight into the distribution and bycatch of diadromous fish at sea.

Despite diadromous fish vulnerability, targeted fishing within estuaries for numerous species remains (Aprahamian and Walker, 2008; Castelnaud, 2000; Feunteun, 2002; Kappel, 2005; Stratoudakis et al., 2020). Given the very coastal distribution of these species, more detailed analysis into such catch data (small fishing vessels targeting diadromous fish) is required. Smaller vessels and artisanal fisheries are still largely overlooked, regardless of their potential threat to diadromous fish (Beaulaton et al., 2008; Castelnaud, 2000; Stratoudakis et al., 2016). Having a better understanding of fishing pressure, combined with the outputs of the distributions and gear capture from these models, could help improve understanding of fishing impacts on diadromous fish.

Finally, it has been well acknowledged that although a large impetus for the designation of MPAs has been undertaken in the last two decades (Probst et al., 2021; Worm et al., 2006). The implementation of management measures has been slow to put in place (Dureuil et al., 2018; Probst et al., 2021; Stratoudakis et al., 2016). Furthermore, little has been undertaken to protect diadromous fish within marine ecosystems (Dureuil et al., 2018; Probst et al., 2021; Stratoudakis et al., 2016). We highlight the value of Habitat Directive MPAs that have the jurisdiction to protect diadromous fish. Limiting access of fisheries with a higher probability of capture to such areas could provide additional protection for diadromous fish and their habitats.

## Acknowledgements

We thank the funders of the project Management of Diadromous Fish in their Environment, OFB-INRAE-Institute Agro-UPPA. We are extremely grateful to all those who were involved in collecting and compiling the fisheries-dependent and fisheries-independent data and their funders. We are also grateful to IFREMER and the French marine fisheries and aquaculture administration (DPMA) for access to their data. Finally, thanks the reviewers for their valuable contribution.

