## Supplementary material for "Modelling the distribution of rare and data-poor diadromous fish at sea for protected area management" (from another version of this preprint: version 1, 2022-10-25)

### Appendix A. Supplementary material

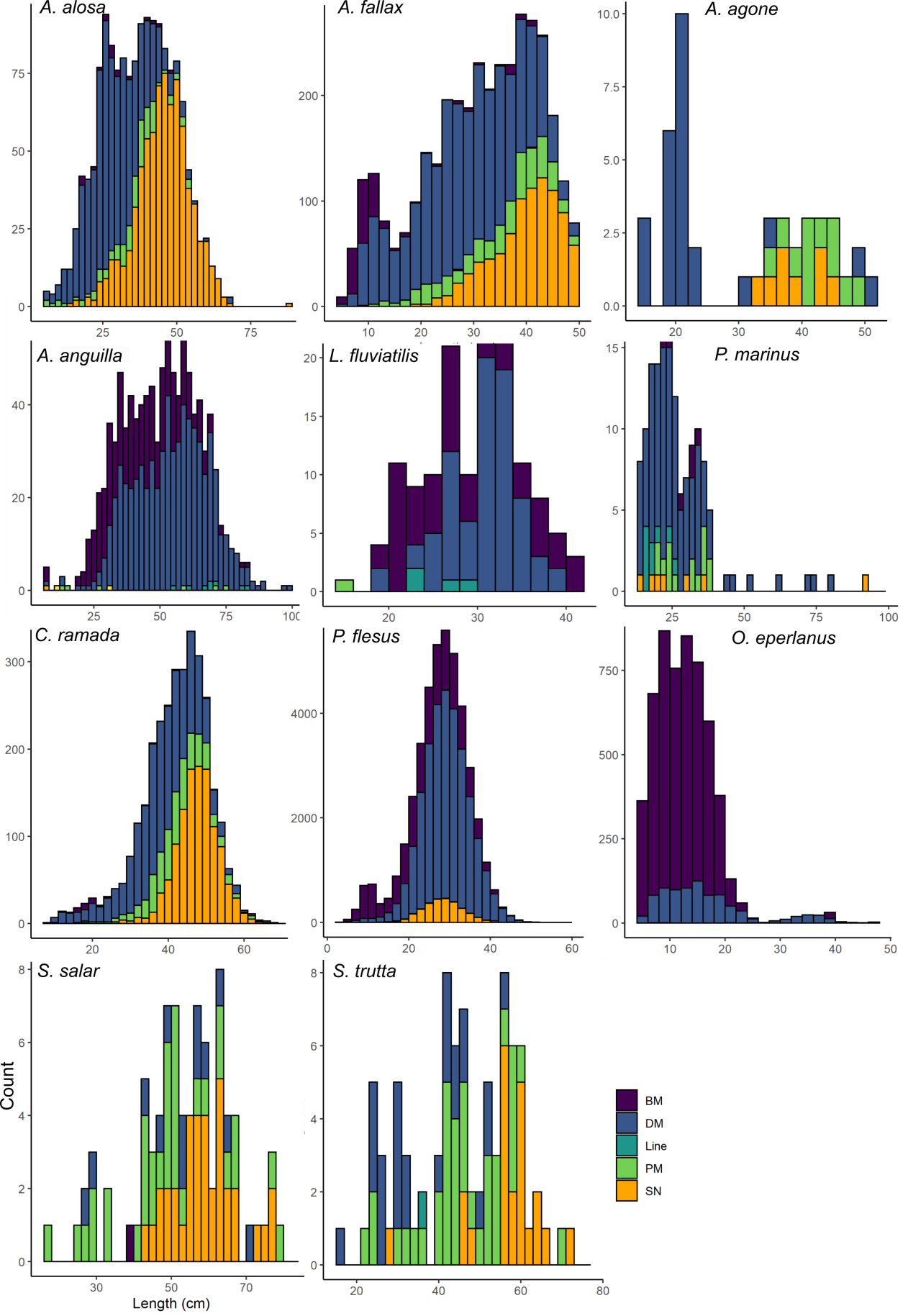

##### **Fig. S1**. Length histogram of diadromous fish caught from the database (2003- 2019) by the different gear categories. BM = Benthic mobile, DM = Demersal mobile, PM = Pelagic mobile, SN = Seine net (refer to Table S3 for gears and their gear category).

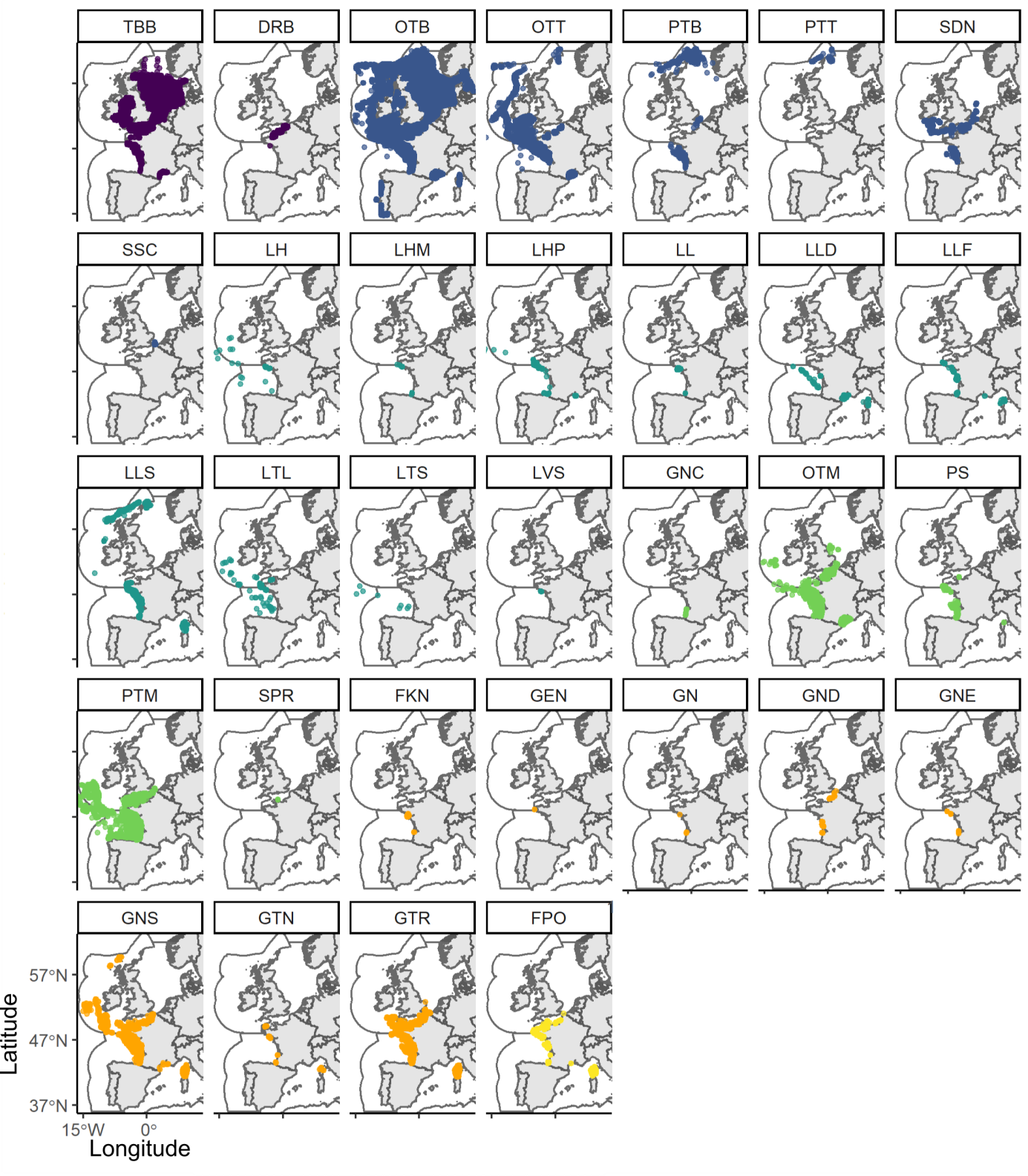

##### **Fig. S2**. Location of the hauls per gear. Grey lines refer to MSFD regions. Colours refer to gear categories: BM = Benthic mobile (purple), DM = Demersal mobile (blue), line (turquoise), PM = Pelagic mobile (green), SN = Seine net (orange), FPO = Pots (yellow; refer to Table S3 for gears and their gear categories).

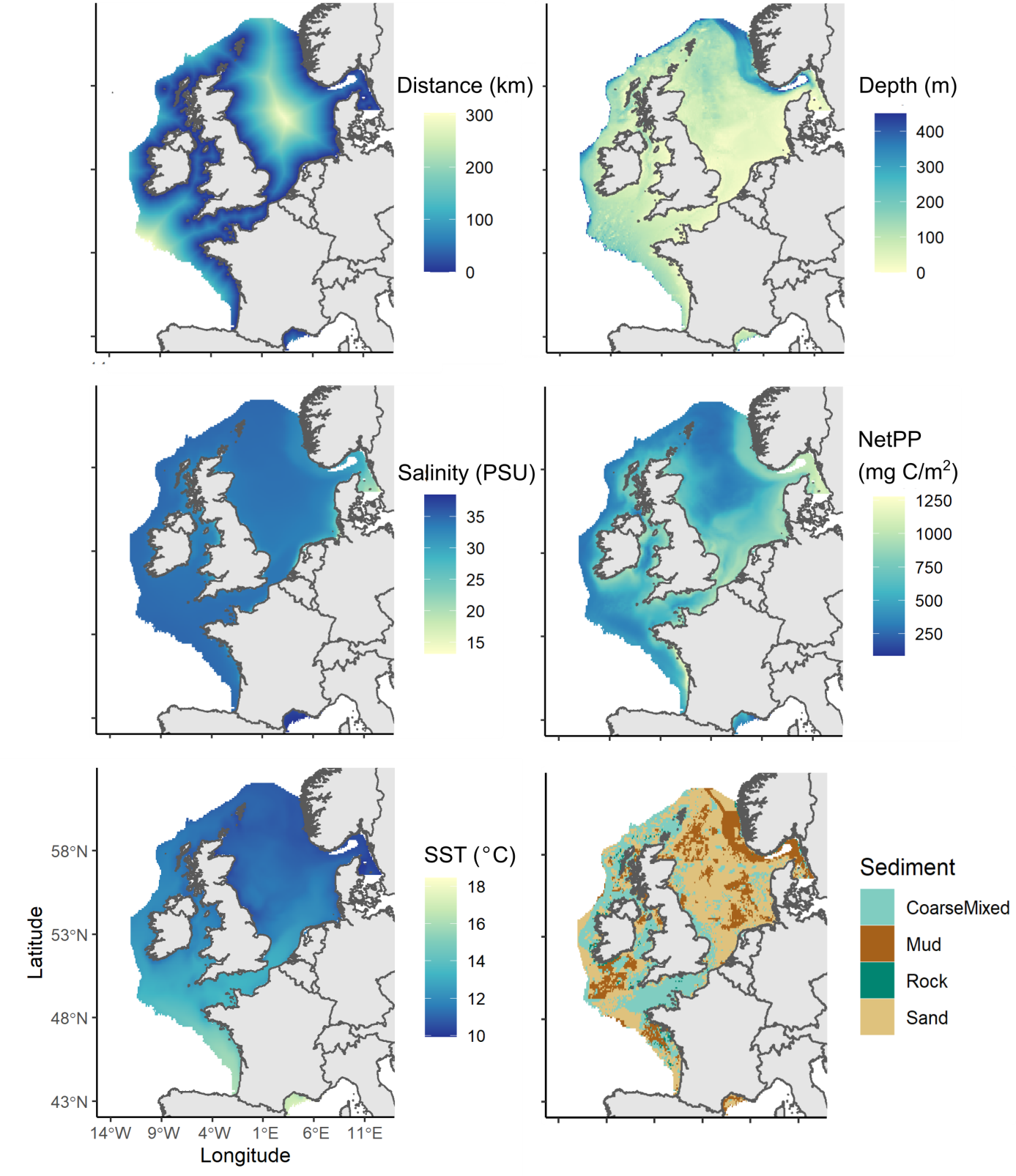

##### **Fig. S3.** Predictor variable maps at a resolution of 0.1 x 0.1 decimal degree grid. SST = Sea surface temperature, NetPP = Net Primary Production:

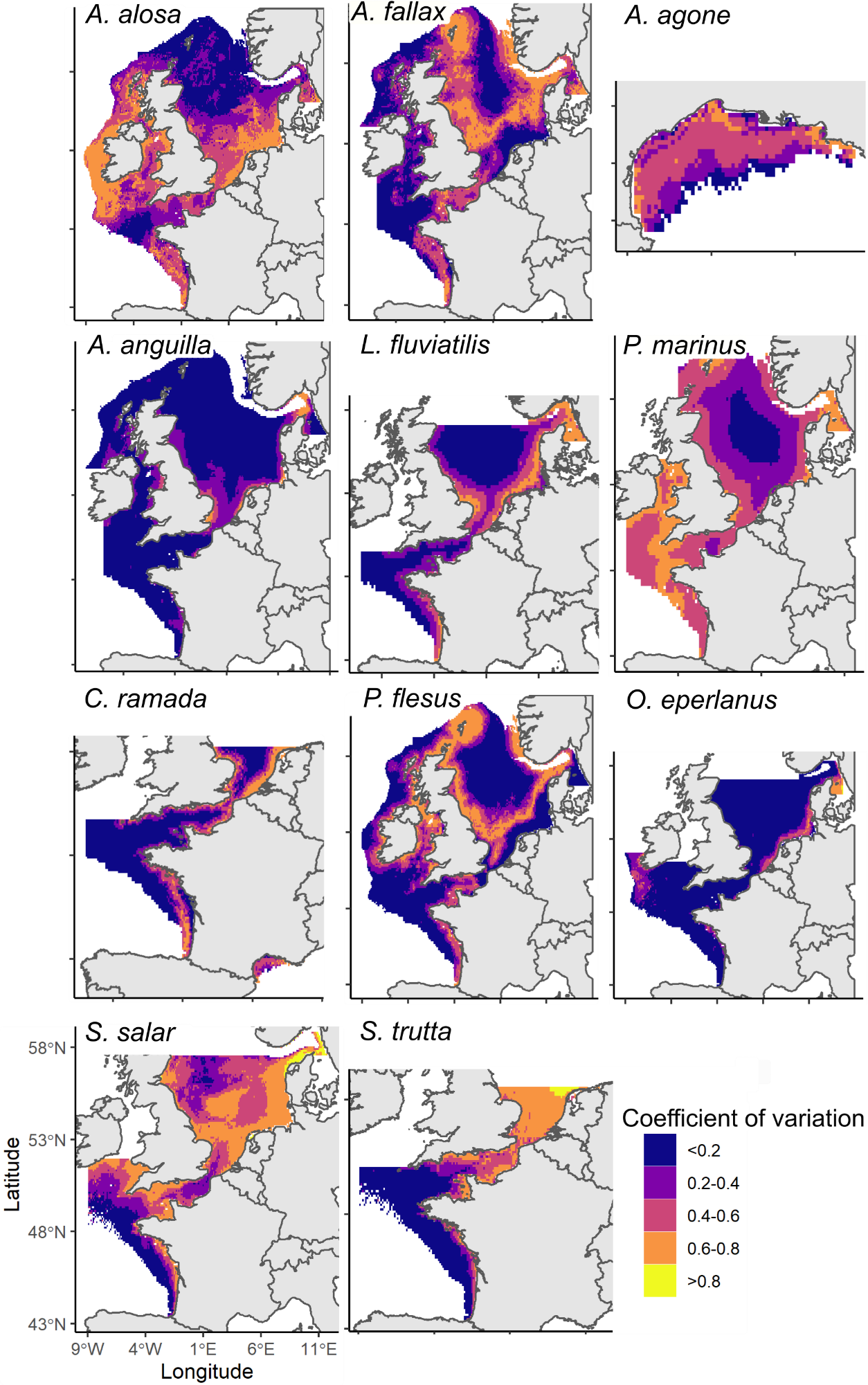

##### **Fig. S4**. Hierarchical species distribution uncertainty maps (calculated by the mean coefficient of variation from the posterior distribution of spatial parameters in the model of best fit).

##### **Table S1.** Diadromous fish hierarchical species distribution model of best fit actual and proportion confusion matrices.

|  |  | **Actual** |  | **Proportion** |  |
| --- | --- | --- | --- | --- | --- |
| **Species** | | **Observed** | |  |  |
|  | **Predicted** | 1 | 0 | 1 | 0 |
| *A. alosa* | 1 | 283 | 454 | 0.04 | 0.07 |
|  | 0 | 75 | 5591 | 0.01 | 0.87 |
| *A. fallax* | 1 | 586 | 1176 | 0.09 | 0.17 |
|  | 0 | 78 | 4963 | 0.01 | 0.73 |
| *A. agone* | 1 | 81 | 164 | 0.11 | 0.22 |
|  | 0 | 19 | 472 | 0.03 | 0.64 |
| *A. anguilla* | 1 | 49 | 50 | 0.01 | 0.01 |
|  | 0 | 44 | 5303 | 0.01 | 0.97 |
| *P. marinus* | 1 | 34 | 38 | 0.02 | 0.02 |
|  | 0 | 23 | 1775 | 0.01 | 0.95 |
| *L. fluviatilis* | 1 | 34 | 57 | 0.03 | 0.05 |
|  | 0 | 14 | 1071 | 0.01 | 0.91 |
| *O. eperlanus* | 1 | 140 | 40 | 0.03 | 0.01 |
|  | 0 | 33 | 4907 | 0.01 | 0.96 |
| *C. ramada* | 1 | 252 | 165 | 0.09 | 0.06 |
|  | 0 | 48 | 2305 | 0.02 | 0.83 |
| *P. flesus* | 1 | 1004 | 408 | 0.14 | 0.06 |
|  | 0 | 109 | 5661 | 0.02 | 0.79 |
| *S. salmo* | 1 | 30 | 85 | 0.01 | 0.02 |
|  | 0 | 20 | 3732 | 0.01 | 0.97 |
| *S. trutta* | 1 | 32 | 120 | 0.02 | 0.06 |
|  | 0 | 13 | 1764 | 0.01 | 0.91 |

##### **Table S2.** Diadromous fish presence per gear type used within the hierarchical species distribution model. Refer to Table S3 for gears acronyms. BM = Benthic Mobile, DM = Demersal Mobile, PM = Pelagic Mobile and SN = Static Net.

| **Latin name** | **Gear category** | **Gear types** | **Number of hauls** | **Number of presences** | **Percent presence (%)** |
| --- | --- | --- | --- | --- | --- |
| *A. alosa* | SN | GND | 60 | 5 | **8.33** |
|  | SN | GNE | 39 | 2 | 5.13 |
|  | SN | GNS | 6674 | **242** | 3.63 |
|  | DM | GTR | 16 516 | 214 | 1.30 |
|  | PM | OTB | 38 060 | **217** | 0.57 |
|  | DM | OTM | 1593 | 23 | 1.44 |
|  | DM | OTT | 15 732 | 26 | 0.16 |
|  | PM | PTB | 1343 | 1 | 0.07 |
|  | DM | PTM | 2882 | 35 | 1.21 |
|  | DM | SDN | 2740 | 26 | 0.95 |
|  | DM | SSC | 28 | 5 | **17.86** |
|  | BM | TBB | 623 | 6 | 0.96 |
| *A. fallax* | SN | GND | 25 | 2 | **8.00** |
|  | SN | GNE | 15 | 2 | **13.33** |
|  | SN | GNS | 5456 | 109 | 2.00 |
|  | SN | GTR | 15 838 | **205** | 1.29 |
|  | Line | LLS | 2175 | 2 | 0.09 |
|  | DM | OTB | 34 890 | **689** | 1.97 |
|  | PM | OTM | 1601 | 25 | 1.56 |
|  | DM | OTT | 14 422 | 62 | 0.43 |
|  | DM | PTB | 320 | 9 | 2.81 |
|  | PM | PTM | 2954 | 112 | 3.79 |
|  | DM | SDN | 2711 | 47 | 1.73 |
|  | BM | TBB | 10 728 | 121 | 1.13 |
| *A. agone* | SN | GNS | 8 | 3 | **37.50** |
|  | DM | OTB | 2680 | **145** | 5.41 |
|  | PM | OTM | 1586 | **17** | 1.07 |
|  | DM | OTT | 816 | 11 | 1.35 |
| *A. anguilla* | SN | GNS | 1789 | 1 | 0.06 |
|  | SN | GTR | 12 466 | 1 | 0.01 |
|  | Line | LLS | 504 | 1 | 0.20 |
|  | DM | OTB | 27 305 | **139** | **0.50** |
|  | PM | OTM | 1435 | 2 | 0.14 |
|  | BM | TBB | 11 055 | **32** | **0.29** |
| *L. fluviatilis* | Line | LLS | 93 | 1 | **1.07** |
|  | DM | OTB | 8216 | **26** | 0.32 |
|  | PM | PTM | 20 | 1 | **0.50** |
|  | DM | TBB | 8300 | **41** | 0.49 |
| *P. marinus* | SN | GNS | 1593 | 4 | 0.25 |
|  | SN | GTR | 6169 | 3 | 0.05 |
|  | Line | LLS | 539 | 8 | **1.48** |
|  | DM | OTB | 15 479 | **35** | 0.23 |
|  | PM | OTM | 909 | 6 | 0.66 |
|  | DM | PTB | 23 | 1 | **4.35** |
|  | PM | PTM | 844 | 8 | 0.95 |
|  | BM | TBB | 3026 | **9** | 0.30 |
| *C. ramada* | PM | GNC | 21 | 1 | **4.80** |
|  | SN | GND | 25 | 2 | **8.00** |
|  | SN | GNS | 5678 | 115 | 2.05 |
|  | SN | GTR | 12 911 | **276** | 2.14 |
|  | DM | OTB | 20 895 | **294** | 1.41 |
|  | PM | OTM | 3187 | 71 | 2.22 |
|  | DM | OTT | 3281 | 12 | 0.37 |
|  | PM | PS | 216 | 6 | 2.78 |
|  | DM | PTB | 124 | 3 | 2.42 |
|  | PM | PTM | 3011 | 95 | 3.15 |
|  | DM | SDN | 2195 | 45 | 2.05 |
|  | PM | TBB | 1539 | 6 | 0.32 |
| *P. flesus* | BM | DRB | 375 | 17 | 4.53 |
|  | SN | GNS | 4520 | 8 | 0.18 |
|  | SN | GTR | 15 683 | 911 | **5.80** |
|  | Line | LLS | 1884 | 1 | 0.05 |
|  | DM | OTB | 38 126 | **2178** | 5.70 |
|  | PM | OTM | 627 | 13 | 2.07 |
|  | DM | OTT | 14 951 | 53 | 0.35 |
|  | PM | PTM | 1761 | 1 | 0.57 |
|  | DM | PTT | 45 | 1 | 2.22 |
|  | DM | SDN | 2706 | 37 | 1.37 |
|  | BM | TBB | 11 722 | **2174** | **18.55** |
| *O. eperlanus* | DM | OTB | 23 709 | **77** | **0.32** |
|  | DM | OTT | 13 855 | 18 | 0.13 |
|  | DM | SDN | 1877 | 1 | 0.05 |
|  | BM | TBB | 9436 | **939** | **9.95** |
| *S. salar* | SN | GNE | 18 | 3 | **16.7** |
|  | SN | GNS | 2302 | **21** | 0.91 |
|  | SN | GTR | 11 003 | 3 | 0.03 |
|  | DM | OTB | 14 165 | 11 | 0.07 |
|  | PM | OTM | 1367 | 6 | 0.44 |
|  | DM | OTT | 1905 | 1 | 0.05 |
|  | PM | PTM | 2358 | **22** | **0.93** |
|  | BM | TBB | 2890 | 1 | 0.03 |
| *S. trutta* | SN | GNS | 2770 | **17** | 0.61 |
|  | SN | GTR | 12 795 | 3 | 0.02 |
|  | Line | LLS | 822 | 1 | 0.12 |
|  | DM | OTB | 11 421 | **19** | 0.17 |
|  | PM | OTM | 603 | 7 | **1.16** |
|  | DM | OTT | 1826 | 1 | 0.05 |
|  | PM | PTM | 1535 | 15 | **0.98** |

##### **Table S3.** Gear types, acronyms, and their gear category. For more detail refer to Table S4 and Elliott et al., (2021) supplementary material.

| **Gear category** | **Gear acronym** | **Gear type** |
| --- | --- | --- |
| Benthic mobile | DRB | Boat dredges |
| Pots | FPO | Pots |
| Static net | GEN | Gillnet and entangling net |
| Static net | GN | Gillnet |
| Pelagic mobile | GNC | Encircling gillnet |
| Static net | GND | Driftnet |
| Static net | GNE | Set gillnet |
| Static net | GNS | Set gillnet (anchored) |
| Static net | GTN | Combined gillnets-trammel nets |
| Static net | GTR | Trammel net |
| Line | LH | Hand line |
| Line | LHM | Han and pole lines (mechanised) |
| Line | LHP | Hand lines and pole-lines (hand operated) |
| Line | LL | Longlines |
| Line | LLD | Drift longline |
| Line | LLF | Fixed floating longline |
| Line | LLS | Set longline |
| Line | LTL | Trolling lines |
| Line | LTS | Surface longline |
| Line | LVS | Vertical longline |
| Demersal mobile | OTB | Otter beam trawl |
| Pelagic mobile | OTM | Otter midwater trawl |
| Demersal mobile | OTT | Otter twin trawl |
| Pelagic mobile | PS | Purse seine |
| Demersal mobile | PTB | Bottom pair trawl |
| Pelagic mobile | PTM | Midwater pair trawl |
| Demersal mobile | PTT | Demersal pair trawl |
| Demersal mobile | SDN | Danish seine net |
| Pelagic mobile | SPR | Vessel pair seine |
| Demersal mobile | SSC | Scottish seine net |
| Benthic mobile | TBB | Bottom beam trawl |

##### **Table S4.** Surveys collated to assess diadromous fish presence (Updated from Elliott et al, (2021)). Gear acronyms within Table S3. MIK = Midwater Ringnet, FKN = Fyke net. NA = No information given.

| **Survey category** | **Full name (where available)** | **Survey acronym** | **Gear acronym** | **Geographical region** | **Years** | **Source** |
| --- | --- | --- | --- | --- | --- | --- |
| ICES DATRAS^[1]^ | Bottom Trawl Survey | BTS | TBB | Greater North Sea and Celtic Sea | 1987 - 2018 | DOI:10.7489/1967-1 |
|  | Bottom Trawl Survey | BTS-VIII | TBB | Bay of Biscay | 2011-2017 | http://datras.ices.dk |
|  | Demersal Young Fish Survey | DYFS | TBB | Greater North Sea | 2002-2017 | http://datras.ices.dk |
|  | Evaluation Halieutique Ouest de l’Europe | EVHOE | OTB | Celtic sea and Bay of Biscay | 1997-2017 | 10.7489/1958-1 Not |
|  | French Channel Ground Fish Survey | FR-CGFS | OTB | Eastern English Channel | 1988-2018 | DOI: 10.18142/11 |
|  | North Sea International Bottom Trawl Survey | NS-IBTS | OTB | Greater North Sea | 1965-2018 | DOI: 10.7489/1922-1 |
|  | Portuguese International Bottom Trawl Survey | PT-IBTS | OTB | Bay of Biscay and the Iberian coast | 2002-2017 | DOI: 10.7489/1963-1 |
|  | Scottish West Coast Bottom Trawl Survey | SWC-IBTS | OTB | Celtic sea | 1985-2010 | DOI: 10.7489/1957-1 DOI: 10.7489/1924-1 |
| French scientific surveys |  | Arca | TBB | Bay of Biscay | 1995, 1996 | Ifremer |
|  |  | Arca | GN | Bay of Biscay | 1996 |  |
|  |  | Arca | FKN | Bay of Biscay | 1995 |  |
|  | Nourriceries démersaux | Nourdem Bargip/ | OTB | Channel and Bay of Biscay (Loire and Seine) | 2015-2018 | DOI: 10.18142/315 |
|  | Etude de l'impact des marées noires de l'Erika | Black | OTB | Bay of Biscay | 2000 | DOI: 10.17600/50150 |
|  |  | Black | TBB | Bay of Biscay | 2000 |  |
|  | Nurse (nourriceries) sur la Baie du mont Saint-Michel | BMSM2003 | TBB | Bay of Biscay | 2003 | Ifremer |
|  | Nurse (nourriceries) sur la baie de Bourgneuf | Bour | TBB | Bay of Biscay | 1980-1984 | Ifremer |
|  | CAmpagne MANche OCcidentale pluridisciplinaire | CAMANOC | OTB | English Channel | 2014 | DOI: 10.17600/14001900 |
|  | Campagne de comparaison de racasseurs lourds et legers sur la sélectivité des soles | Captusol01 | TBB | Bay of Biscay | 1984 | Ifremer |
|  | CLASS 2008 | CLA08_ | OTM | Bay of Biscay | 2008 | DOI: 10.17600/8040050 |
|  |  |  | MIK | Bay of Biscay | 2008 |  |
|  | Analyse des peuplements de crépidules en baie de bourgneuf | CREBOUR | TBB | Bay of Biscay | 2006 | DOI: 10.17600/6050020 |
|  | EPIfaune BENthique du GOlfe du Lion | EPIBENGOL | OTB | Mediterranean | 2018 | DOI: 10.17600/18000589 |
|  |  | Giro | TBB | Western English Channel | 1997 | Ifremer |
|  | Campagne sur les gradients de production côte-large incluant la baie de Vilaine et la rade de Brest | Isobaie09 | OTB | Bay of Biscay | 2009 | Ifremer |
|  |  | Isobaie09 | FKN | Bay of Biscay | 2009 |  |
|  |  | Isobaie09 | NA | Bay of Biscay | 2009 |  |
|  | Nurse (nourriceries) en estuaire externe de la Loire | Loir | TBB | Bay of Biscay | 1980-1986 | Ifremer |
|  |  | Marika | TBB | Bay of Biscay | 2005 | Ifremer |
|  | Mediterranean trawl survey - Corse | MEDITS-CORSE | OTB | Mediterranean | 1994-2017 | http://www.ifremer.fr/SIH-indices-campagnes/ |
|  | Mediterranean trawl survey – Golfe du Lion | MEDITS-LION | OTB | Mediterranean | 1994-2018 | http://www.ifremer.fr/SIH-indices-campagnes/ |
|  | Nurse (nourriceries) en 2003 | MISOLRE | OTB | Bay of Biscay | 2003 | DOI:10.17600/3050120 |
|  |  | MISOLRE | TBB | Bay of Biscay | 2003 |  |
|  | Nurse (nourriceries) dans le golfe du Morbihan | Morb | TBB | Bay of Biscay | 1995-1996 | Ifremer |
|  | Nurse (nourriceries) dans le pertuis breton | Nonamebreton | TBB | Bay of Biscay | 1987 | Ifremer |
|  | Nurse (nourriceries) dans la Vilaine | Nonamvillaine | TBB | Bay of Biscay | 1996 | Ifremer |
|  | Nourriceries en Méditerranée | NOURMED | TBB | Mediterranean | 2018 | DOI: 10.17600/18000588 |
|  | Nourriceries Baie de Somme | NOURSOM | TBB | Channel | 1987-2018 | DOI: [10.18142/22/](https://sextant.ifremer.fr/record/98ac17c5-8d70-455e-9496-aafb0dd4ae17/) |
|  | Nourriceries golfe de Gascogne | Nurse | TBB | Bay of Biscay | 1981-2016 | Ifremer |
|  |  | Nurse | OTB | Bay of Biscay | 2000-2002,  2008-2009 |  |
|  |  | Nurse | NA | Bay of Biscay | 2009 |  |
|  |  | Nurse | GN | Bay of Biscay | 2013 |  |
|  | Nurse (nourriceries) dans la Vilaine | Nurvil | TBB | Bay of Biscay | 2004-2005 | Ifremer |
|  |  | Nurvil | OTB | Bay of Biscay | 2005 |  |
|  |  | Peco/PECOS | OTB | Bay of Biscay and Western English Channel | 1976-1980  2004-2006 | Ifremer |
|  |  | Peco/PECOS | TBB | Bay of Biscay and Western English Channel | 1980, 2004 |  |
|  | Pélagiques Gascogne | PELGAS | OTM | Bay of Biscay | 2000-2018 | DOI: 10.17882/59809 |
|  |  | PELGAS | PS | Bay of Biscay | 2007-2011 |  |
|  |  | PELGAS | PTM | Bay of Biscay | 2007-2018 |  |
|  | Nurse (nourriceries) dans les pertuis (breton et antioche) | Pertuis | TBB | Bay of Biscay | 1986-1987 | Ifremer |
|  | Campagnes pour la compréhension des communautés sur les habitats à Haploops | Ploops | TBB | Bay of Biscay | 2009-2010 | Ifremer |
|  |  | RESSGASC | OTB | Bay of Biscay | 1984-2002 | Ifremer |
|  | Nurse (nourriceries) en baie de Vilaine mais en hiver (winter) | RetroB | TBB | Bay of Biscay | 2008 | Ifremer |
|  | Suivi des Effets bioLogIques de la contamination chimique en Méditerranée | SELIMED-1 | OTB | Mediterranean | 2016 | DOI: 10.17600/16011900 |
|  | Répartition de la distribution des soles sur la radiale cote-large de la baie de Vilaine | Sold | OTB | Bay of Biscay | 1988-1989 | DOI:10.17600/88011211 |
|  |  | Typo | OTB | Bay of Biscay | 1996 | Ifremer |
|  |  | Typo | TBB | Bay of Biscay | 1997-1997 |  |
| Fisheries-dependent data | French fisheries observer data | ObsMer | Refer to Table S3 | Greater North Sea, Celtic Sea, Bay of Biscay | 2003-2019 | DOI: 10.13155/39722 |
